# What’s in Cow Dung Manure? Insights into Its Potential for Supporting Plant Defense System

**DOI:** 10.1101/2024.11.20.624574

**Authors:** Karun Wilson, Sathiavelu Arunachalam

## Abstract

*Solanum lycopersicum*, a prominent vegetable crop, suffers major yield losses from early blight disease driven by *Alternaria solani.* Sustainable alternatives to conventional chemical fungicides are urgently needed to mitigate these losses while minimizing environmental impact. This study investigates the application of the microbial consortium derived from Gir cow dung for enhancing tomato resistance against early blight. Metagenomic analysis of Gir cow dung revealed a rich diversity of beneficial microbes, with *Bacillus* species being one of the prominent. Three key strains *Bacillus subtilis*, *Bacillus tequilensis*, and *Bacillus licheniformis* were isolated, characterized, and tested for their antagonistic activity against *A. solani*. The microbial consortium exhibited significant antifungal properties, and the application of the consortium on tomato plants led to a marked reduction in disease progression. Histochemical analyses of treated plants revealed a pronounced increase of hydrogen peroxide, phenolic compounds, and superoxide dismutase, suggesting enhanced activation of plant defense pathways. Comparative gene expression profiling showed the upregulation of key defense-related genes, including PR2b, Chi3, and Pto kinase, in Bo-treated plants, indicating a systemic activation of host defense mechanisms.

Untargeted metabolomic analysis using LC-MS/MS further demonstrated significant metabolic reprogramming in microbial consortium-treated plants. Pathway enrichment analysis revealed enhanced accumulation of secondary metabolites, including phenolics and terpenoids, which are known to play vital roles in pathogen defense. Additionally, hierarchical clustering of metabolite profiles highlighted distinct metabolic signatures between Bo-treated and control plants, underscoring the broad biochemical impact of the microbial treatment. Collectively, our findings demonstrate that the microbial consortium derived from Gir cow dung significantly enhances tomato plant resistance to *A. solani* through multifaceted biochemical, molecular, and metabolic mechanisms. This study provides a sustainable biocontrol strategy that could effectively replace chemical fungicides, contributing to resilient tomato production and advancing the field of agroecological plant disease management.

## Introduction

*Solanum lycopersicum* is a prominent agricultural crop globally, valued for its nutritional content and economic significance. However, its productivity is frequently challenged by *Alternaria solani* which causes early blight disease (Chaerani and Voorrips 2006; Agrios 2004). Conventional control measures primarily involve chemical fungicides, which, while effective, come with several drawbacks, including the emergence of resistant strains, environmental toxicity, and health hazards for farmworkers and consumers (Brent and Hollomon 2007; Gisi and Sierotzki 2008). Thus, there is an alarming demand for sustainable alternatives to mitigate this pathogen effectively.

Using beneficial microorganisms in crop development has emerged as a promising strategy for controlling phytopathogens. Among these, members of the *Bacillus* genus are particularly recognized for their plant growth development and biological control properties (Köhl, Kolnaar, and Ravensberg 2019). *Bacillus* species like *Bacillus subtilis*, *Bacillus tequilensis*, and *Bacillus licheniformis* are well-documented for their potential to produce biologically active compounds, counting lipopeptides, antibiotics, and enzymes that inhibit fungal pathogens (Stein 2005; Ongena and Jacques 2008). These bacteria also kick the systemic tolerance in host plants, effectively enhancing their defense responses to subsequent pathogen challenges (Choudhary and Johri 2009). Given their robustness, spore-forming ability, and diverse metabolic capabilities, *Bacillus* species represent an attractive alternative to chemical fungicides for enhancing crop resilience (Kloepper, Ryu, and Zhang 2007).

In the framework of sustainability, the utilization of organic manure, such as cow-dung, shows a dual part in providing essential nutrients and harboring a diverse microbial community beneficial for plant health (S. Zhang et al. 2020). Cow-dung manure is a good source of beneficial microbes (Gupta, Aneja, and Rana 2016), including *Bacillus* species, that can be leveraged for biocontrol and growth promotion (Ji et al. 2022). The potential of microbial consortia derived from cow dung has not only demonstrated efficacy in enhancing plant growth but also in providing a sustainable means of reducing pathogen load through natural microbial antagonism.

To elucidate the mechanisms through which these microbial consortia enhance plant resistance, it is essential to integrate advanced analytical tools, such as metagenomics, untargeted metabolomics, and gene expression profiling. Metagenomic sequencing enables the characterization of the microbial diversity in cow dung and helps in identifying key beneficial microorganisms (Nam et al. 2023). Untargeted metabolomics, using technologies like LC-MS/MS, allows for a comprehensive understanding of the metabolic shifts that occur in tomato plants treated with microbial consortia, particularly focusing on metabolites associated with defense responses and secondary metabolite pathways (Bozza et al. 2024). These metabolic insights, combined with gene expression profiling of critical defense-related genes, such as *PR2b*, *Chi3*, and *Pto* kinase, provide a holistic view of the biochemical and molecular variations encouraged in tomato plants in retort to microbial treatment (Conrath 2006; Pieterse et al. 2014).

In this research, we isolated and identified microbial communities from Gir cow dung to evaluate their potential to enhance tomato plant resistance against early blight caused by *A. solani*. Specifically, we focused on isolating *Bacillus* strains with antifungal activity and assessed their effects on disease progression in tomato plants. Using metagenomics, we characterized the microbial diversity of Gir cow dung, while untargeted metabolomics and gene expression studies were employed to unravel the changes in plant metabolic and defense pathways induced by the microbial consortium. Our results demonstrate that treatment with the microbial consortium significantly reduces disease severity and promotes systemic defense responses, as evidenced by an increased buildup of H₂O₂, (hydrogen peroxide) phenolic compounds, and SOD (superoxide dismutase), also the upregulation of defense-related genes. These findings highlight the potential of microbial consortia as a sustainable biocontrol strategy, reducing dependence on chemical fungicides and promoting resilient agricultural practices (Harman et al. 2004; van Loon 2000).

## Results

### Nutrient Composition of Gir Cow Dung

A comprehensive nutrient analysis of the Gir cow dung sample was conducted to assess its suitability for agricultural application. The sample exhibited (Table 2) a neutral pH of 7.5, indicating that it would not adversely alter soil pH upon application. The moisture content was 34.91%, suggesting the dung’s ability to retain adequate moisture, which is beneficial for microbial activity. The macronutrient profile, determined through standardized protocols, revealed total nitrogen content at 0.80%, total phosphate (P₂O₅) at 1.70%, and total potash (K₂O) at 0.37%. These values indicate that Gir cow-dung is a potential means of essential nutrients required for plant growth. The total organic carbon content was found to be 23.36%, with a carbon-to-nitrogen ratio of 29:1, favoring the growth of beneficial microbes.

**Table 1:**
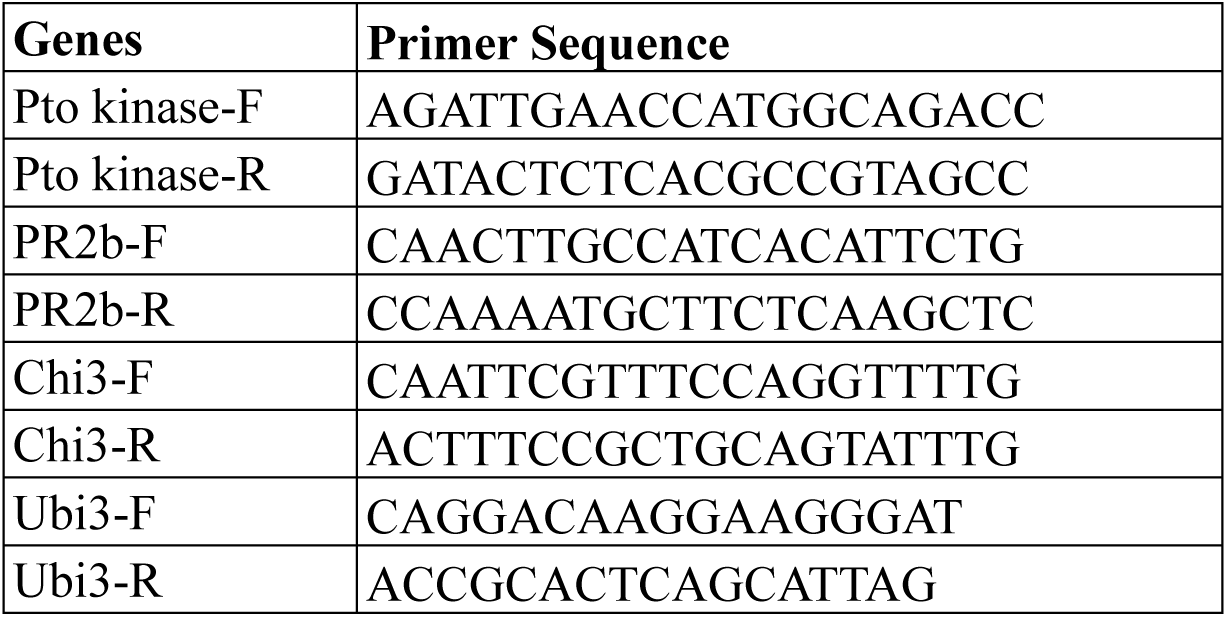
Primer Sequences for Gene-Specific RT-PCR Analysis.

**Table 2:**
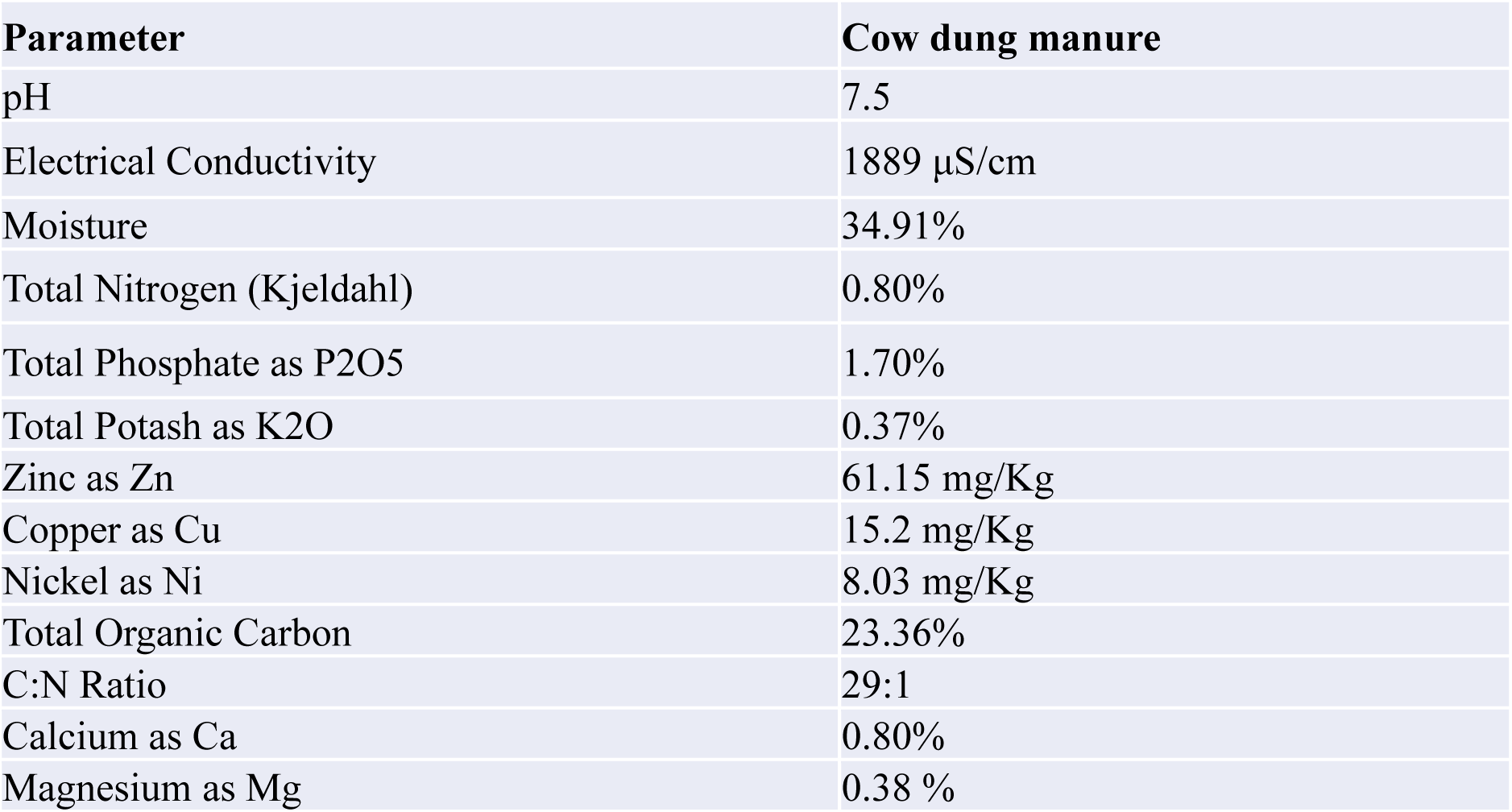
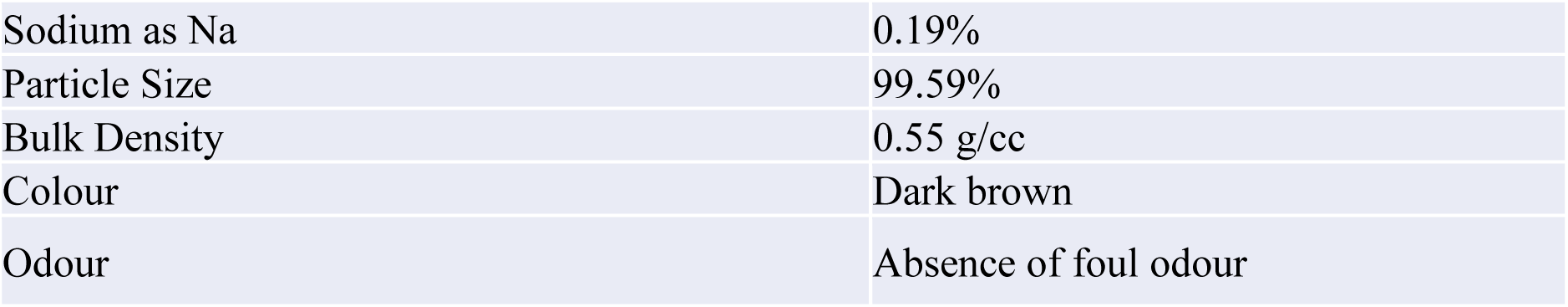
Physicochemical properties and nutrient composition of cow dung manure.

The analysis of trace showed optimum levels of zinc, copper, and nickel, these micronutrients play crucial roles in plant metabolism and can enhance soil fertility when applied as a soil amendment. Additional minerals, including calcium, magnesium, and sodium, were also quantified. The physical characteristics of the cow dung, such as bulk density (0.55 g/cc) and particle size distribution (99.59%), indicate its suitability for improving soil structure. The absence of a foul odor and the dark brown color suggest proper decomposition, making it more acceptable for agricultural use.

Overall, the nutrient profile of Gir cow dung demonstrates its potential as a rich source of macro-and micronutrients, organic matter, and microbial support, which can be utilized for promoting plant growth and enhancing soil health.

### Bacterial diversity in Gir Cow Dung

The 16S rRNA amplicon sequencing of the Gir cow dung sample (CD-Gir) revealed a diverse bacterial community, including various taxa identified for their potential antifungal properties and plant growth-enhancing abilities Fig. 1 (A, B, C).

**Figure 1:**
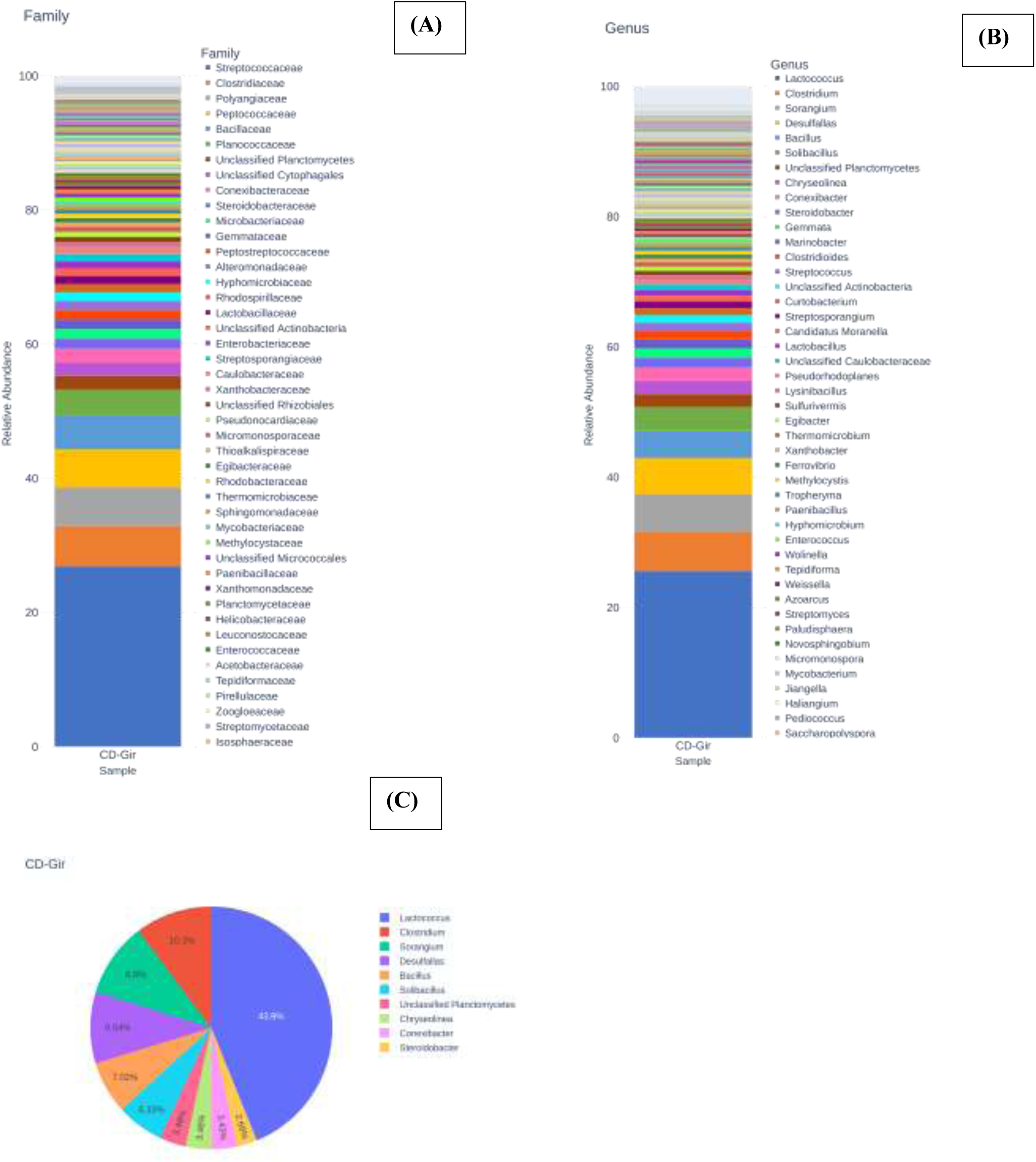
Bacterial Community Composition in Gir Cow Dung Based on 16S rRNA Amplicon Sequencing. The figure 1 represents the taxonomic diversity observed in the microbial community of Gir cow dung at different taxonomic levels. **(A)** A bar plot representing the relative abundance of bacterial families in the sample, providing insight into the broader taxonomic classification and illustrating the variety of bacterial families present. **(B)** The bar plot depicts the relative richness of bacterial genera in the cow dung sample, presenting a detailed breakdown of the diversity beyond the most dominant taxa. **(C)** Pie chart showing the relative richness of dominant bacterial species in the cow dung sample (CD-Gir), highlighting that *Lactococcus* is the most abundant genus, followed by *Clostridium*, *Sorangium*, and others.

**Figure 2:**
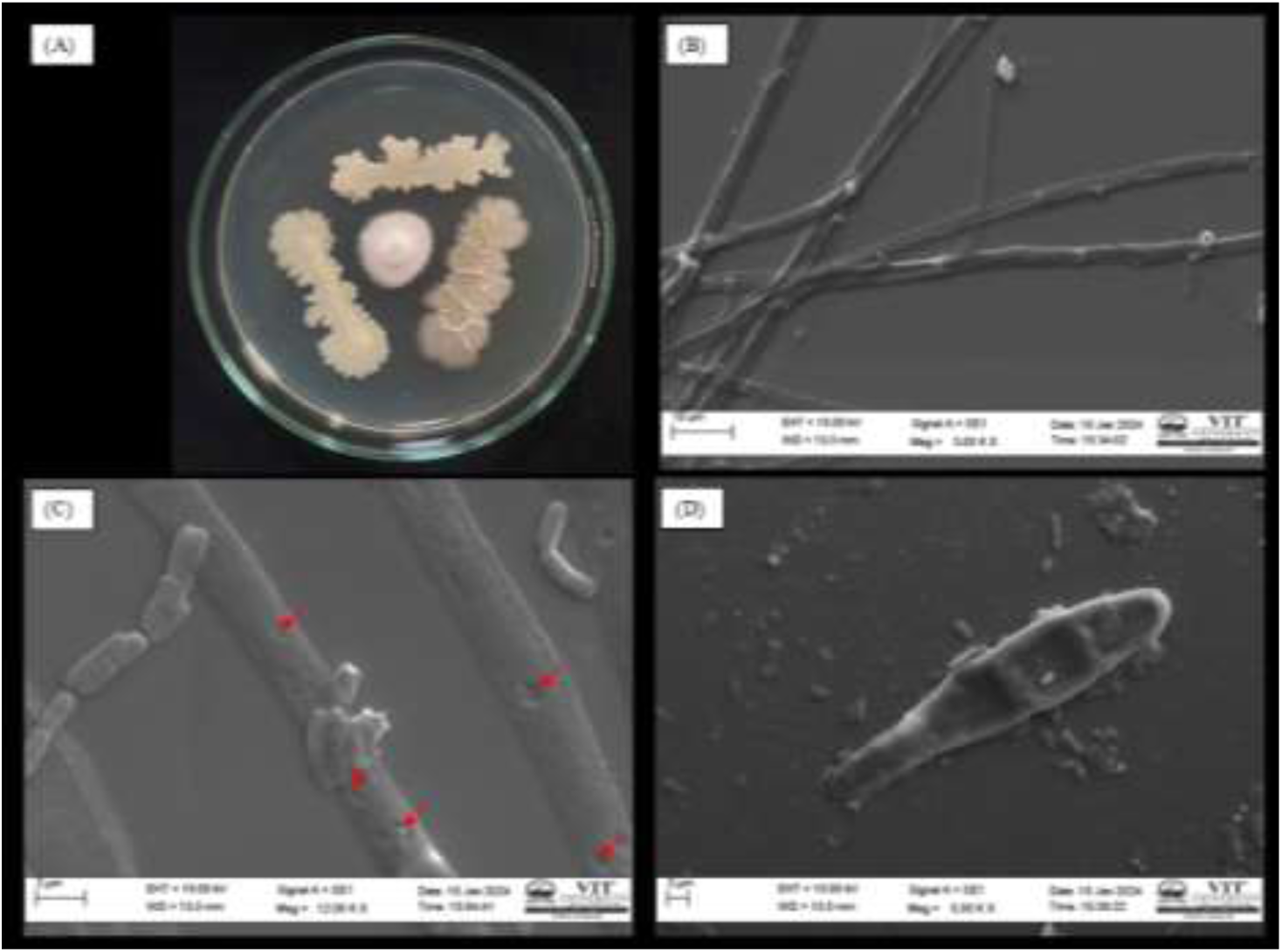
Fungal plate showing the antagonistic activity of *B. licheniformis*, *B. subtilis*, and *B. tequilensis* against *A. solani* on a PDA plate. (B) SEM image of *A. solani* fungal mycelia (control), displaying an intact, smooth surface. (C) SEM image of *A. solani* fungal mycelia interacting with the microbial consortium, with the mycelial surface showing deformation of the hyphal structure (indicated by red arrows). (D) Intact *A. solani* spore interacting with the microbial consortium.

At the family level, the bacterial community was dominated by *Streptococcaceae* and *Bacillaceae* (Figure 1 A). *Streptococcaceae* was the most prevalent family in the sample, followed by *Clostridiaceae* and *Bacillaceae. Bacillaceae* is particularly notable due to its members’ well-documented roles in plant growth promotion and biocontrol activities. Other families, such as *Polyangiaceae, Peptococcaceae*, and *Microbacteriaceae*, were also present in considerable abundance.

At *the* genus level *Lactococcus* was the most abundant, accounting for 43.9% of the total bacterial community (Figures 1B and 1C). *Clostridium* was the second most abundant genus, representing 10.3%, followed by *Sorangium* (9.9%) and *Desulfallas* (9.54%). *Bacillus*, a genus well-known for its antifungal properties and plant growth-promoting activities, constituted 7.02% of the total bacterial community. Additionally, genera such as *Solibacillus* (6.33%) and *Streptomyces* were identified, among which includes species documented for roles in promoting plant health and suppressing pathogens.

### Potential Functional Implications

The dominance of *Lactococcus* and the substantial presence of *Bacillus* within the microbial community suggest a microbiome that could be rich in bacteria with antifungal and plant growth-promoting properties. The diverse composition, including both high-abundance and less abundant taxa known for their beneficial roles, indicates that Gir cow dung harbors a complex microbial community that may contribute to plant health and defense.

### Isolation and Screening of Antagonists

An overall of 53 bacterial species were obtained from the Gir cow dung sample through serial dilution and plating on nutrient agar. The isolates were purified based on distinct colony morphologies by repeated streaking on fresh agar plates. The antagonistic ability of all bacterial species against *A. solani* was evaluated using a dual culture assay. Three isolates demonstrated significant antifungal activity, with inhibition rates exceeding 50%. These isolates were recognized via 16S rRNA gene sequencing as *Bacillus licheniformis*, *Bacillus subtilis*, and *Bacillus tequilensis*. Given their strong antagonistic activity, these strains were chosen for the characterization.

### Secondary metabolite analysis of the bacterial extract by GC–MS

GC-MS analysis of bacterial extracts from *B. subtilis*, *B. tequilensis*, and *B. licheniformis* identified several common compounds, demonstrating a consistent metabolic profile across these strains (Figure 3). The most dominant compound was Acetic acid ethyl ester, with the highest concentration observed in *B. tequilensis* (94.06%), followed by *B. licheniformis* (86.69%) and *B. subtilis* (79.69%). This suggests that ethyl acetate-based compounds are major metabolites in all three *Bacillus* species analyzed.

**Figure 3.**
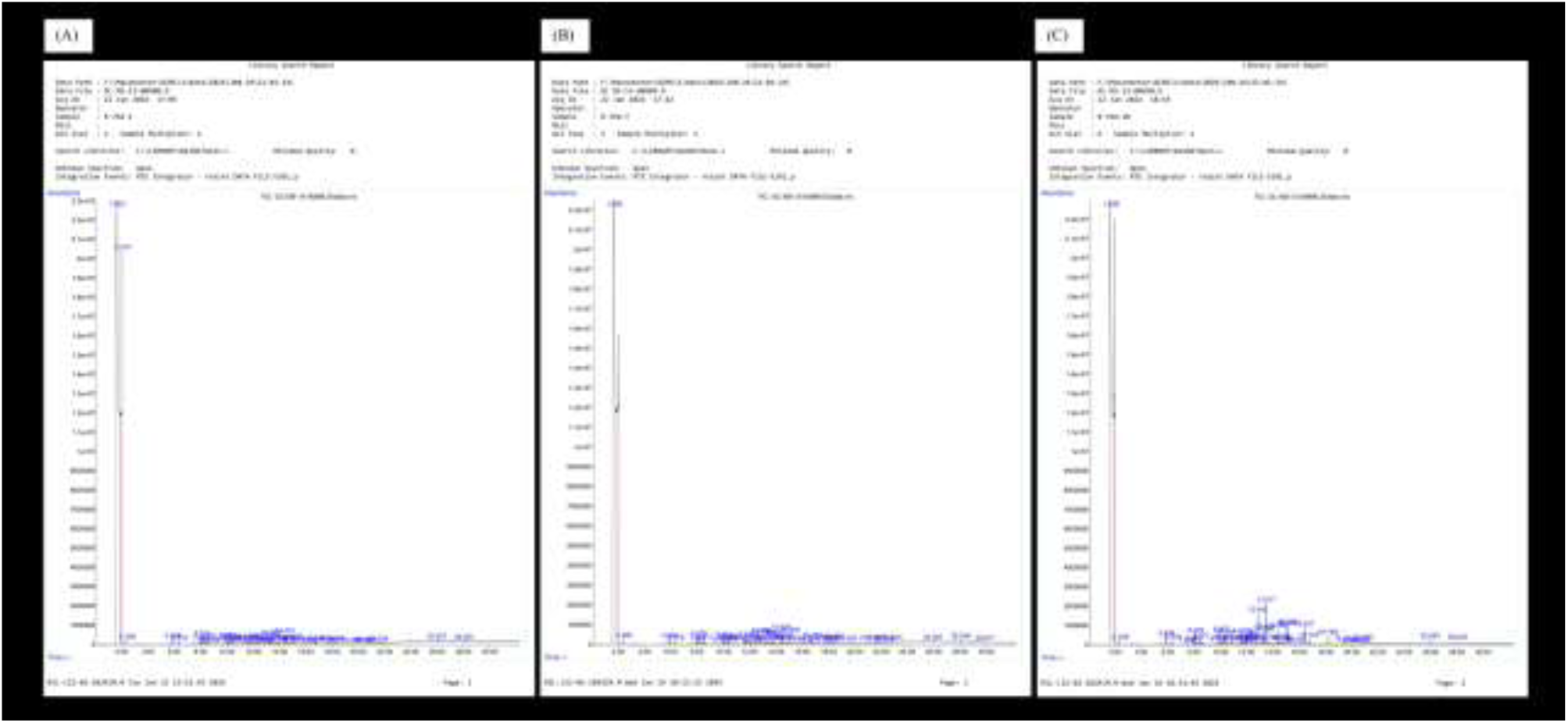
GC-MS chromatograms of ethyl acetate extracts from three bacterial samples. (A) *Bacillus subtilis*, (B) *Bacillus tequilensis*, and (C) *Bacillus licheniformis*. The chromatograms represent the elution profiles of compounds, with retention times (x-axis) and relative richness (y-axis) indicated. Peaks represent bioactive compounds identified in the bacterial extracts, showing distinct profiles for each sample.

Propanoic acid, ethyl ester was another common compound found in all three samples, although at lower concentrations. It was detected at 0.15% in *B. tequilensis*, 0.12% in *B. licheniformis*, and 0.09% in *Bacillus subtilis*. Similarly, Tricosane, a hydrocarbon known for its role in antimicrobial activity, was present in higher amounts in *B. licheniformis* (0.43%) compared to *B. tequilensis* (0.22%) and *B. subtilis* (0.02%).

Additionally, 2,4-Di-tert-butylphenol, a compound recognized for its potent antimicrobial properties, was identified in all three samples, with concentrations of 0.11% in *B. licheniformis*, 0.08% in *B. tequilensis*, and 0.07% in *B. subtilis*. Another notable shared compound, (1R,3S,5S,8R)-3-acetyl-1,5-dimethyl-6-oxabicyclo [3.2.1] octan-7-one, was detected at 0.38% in *B. licheniformis*, 0.30% in *B. tequilensis*, and 0.17% in *B. subtilis*. The detection of bioactive compounds such as 2,4-Di-tert-butylphenol suggests the potential antimicrobial and antifungal activity of these *Bacillus* species, highlighting their possible application in biocontrol and plant health promotion strategies.

### Biochemical Characterization of Antagonistic Bacteria

The biochemical characterization of the three *Bacillus* species was executed using the KB001 kit (Table 3). All isolates tested negative for methyl red, indole production, and Voges-Proskauer tests. In terms of carbohydrate fermentation, *B. subtilis* and *Bacillus licheniformis* showed positive reactions for glucose, arabinose, and mannitol. However, *B. tequilensis* was negative for these tests, except for citrate utilization, where it was the only isolate to test positive. All isolates were negative for the fermentation of adonitol, lactose, sorbitol, and rhamnose. These biochemical profiles support the identification of the isolates and provide insights into their metabolic capabilities.

**Table 3.** Biochemical Characterization of Antagonistic Bacteria Using KB001 Kit.

| No | Test | *Bacillus subtilis* | *Bacillus tequilensis* | *Bacillus licheniformis* |
| --- | --- | --- | --- | --- |
| 1 | Indole | -ve | -ve | -ve |
| 2 | Methyl red | -ve | -ve | -ve |
| 3 | Voges-Proskauer | -ve | -ve | -ve |
| 4 | Citrate utilization | -ve | +ve | -ve |
| 5 | Glucose | -ve | -ve | +ve |
| 6 | Adonitol | -ve | -ve | -ve |
| 7 | Arabinose | +ve | -ve | +ve |
| 8 | Lactose | -ve | -ve | -ve |
| 9 | Sorbitol | -ve | -ve | -ve |
| 10 | Mannitol | +ve | -ve | +ve |
| 11 | Rhamnose | -ve | -ve | -ve |
| 12 | Sucrose | +ve | -ve | -ve |

### Plant Growth-Promoting Properties

All three *Bacillus* isolates exhibited positive responses in the plant growth-promoting assays. Siderophore production was evident in all isolates, as shown by the development of a halo area on chrome azurol S (CAS) agar plates. The isolates also confirmed the capacity to solubilize the phosphate, as shown by the clear halo area formed on Pikovskaya’s agar. Their nitrogen-fixing capability was confirmed by their growth in nitrogen-free Jensen’s medium. Additionally, all isolates produced indole-3-acetic acid (IAA), with the progress of pink color in the presence of Salkowski’s reagent and absorbance readings at 530 nm confirming IAA production. These results indicate that the isolates possess multiple plant growth-promoting properties, along with antifungal activity, highlighting their potential for enhancing plant defense and growth.

### Evaluation of Disease Development in Control (CO) and Treated (BO) Plants

The PKM-1 tomato seedlings were inoculated with spore suspension of *A. solani*, leading to faster disease advancement in the susceptible control (CO) group compared to the resistant consortium treated (BO) group, as shown in Fig. 4 (A). By 96 hours after inoculation, CO plants exhibited pronounced necrotic spots. Disease development was assessed using a detached leaf method. After 24 hours of infection, small brownish spots appeared, gradually enlarging and spreading across the leaf surface. During the first 48 hours, changes in the size of lesions were not substantial in either group. However, by 72 and 96 hours, a clear difference in lesion expansion was observed between Bo and Co plants. At each time point, the spread of the disease in CO plants was notably higher (P < 0.05) than in BO plants. At 96 hours, the largest lesion in CO plants reached about 10.2 mm, whereas in BO plants, it measured only 6.5 mm. Furthermore, the necrotic area on leaves in CO plants was nearly twice as large as that of BO plants, highlighting the greater resistance observed in BO.

**Figure 4:**
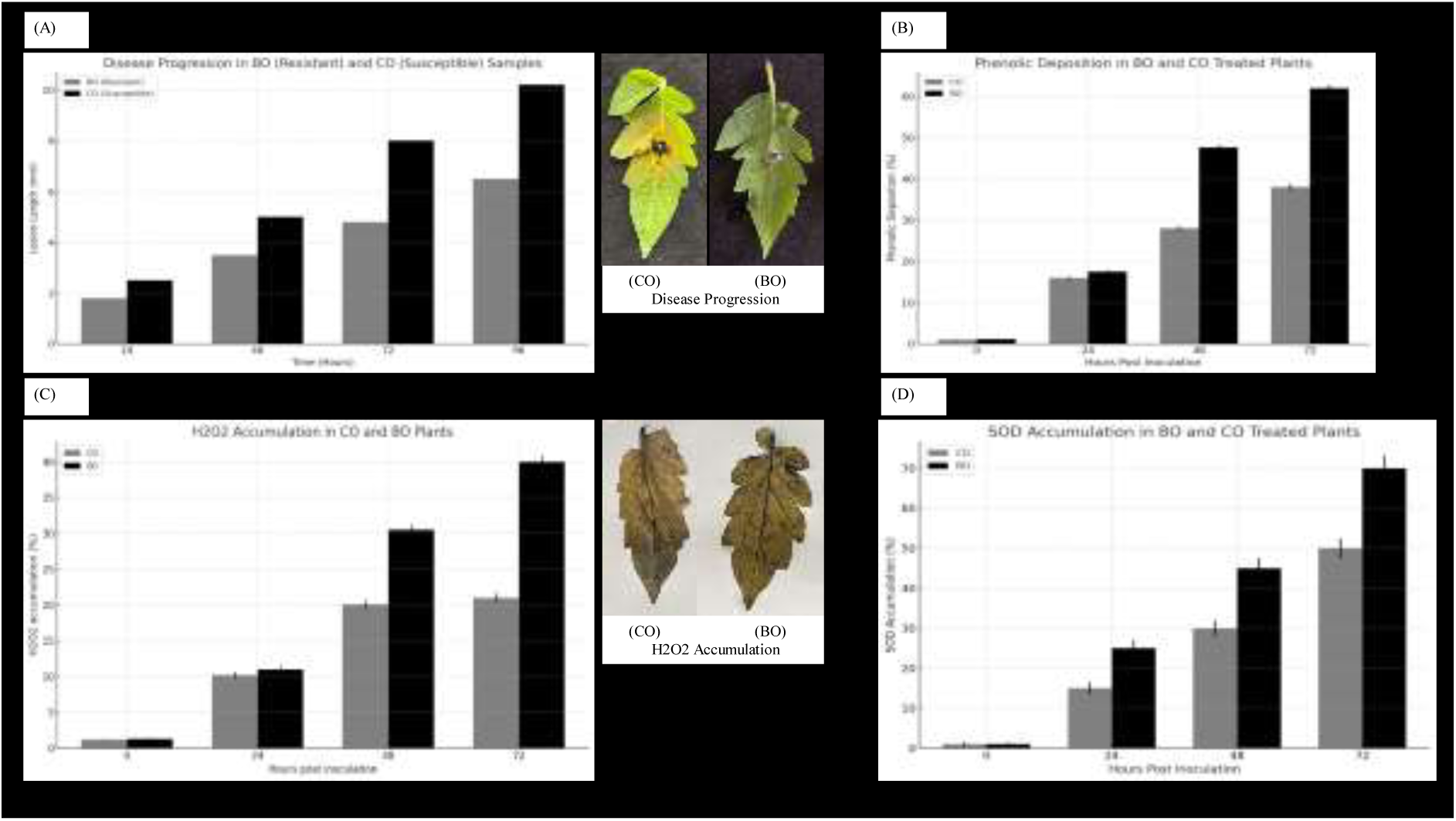
**(A)** Lesion length (mm) in *Alternaria solani*-inoculated tomato leaves of BO (resistant) and CO (susceptible) over 24, 48, 72, and 96 hours. ANOVA with Bonferroni post-hoc test (P < 0.05). Error bars: standard deviation; different letters show significant differences. **(B)** Phenolic compound deposition in primary leaves of BO and CO from 0 to 72 hours. ANOVA with Bonferroni post-hoc test (P < 0.05). Error bars: SEM; different letters denote significant differences. **(C)** H₂O₂ localization in primary leaves of CO and BO from 0 to 72 hours post-inoculation. Independent t-test (P ≤ 0.05). Error bars: SEM; different letters indicate significant differences. **(D)** SOD accumulation in primary leaves of BO (treated) and CO (control) at 0, 24, 48, and 72 hours. ANOVA with Bonferroni post-hoc test (P < 0.05). Error bars: standard deviation; different letters indicate significant differences.

### Localizations of Phenolic Compounds using Histochemical Analysis

The phenolic accumulation in BO-treated and CO-treated samples was assessed to evaluate the reinforcement of the cell wall, a main plant defense mechanism. The phenolic deposition increased steadily in both BO and CO samples, with significant differences emerging at 48 hours post inoculation (Fig. 4 B). At 72 hours, the phenolic accumulation in the BO sample was 62%, significantly higher than the CO sample, which reached 38% (P < 0.05). The BO-treated plants exhibited 1.63-fold higher phenolic deposition than CO-treated plants at 72 hours, indicating enhanced defense responses in BO-treated plants.

### Histochemical accumulation of H₂O₂ comparison in Control (Co) and Treated (Bo) Plants

Histochemical buildup of H2O2 in answer to the pathogen was noted in leaf materials and linked between the control (CO) and BO-treated plants. Both sets of plants showed significant accumulation of H_2_O_2_ starting from 24 hours post-inoculation (P ≤ 0.05). However, BO-treated plants exhibited significantly higher levels of H_2_O_2_ buildup compared to the CO plants at all time points (Fig. 4 C). At 24 hours, the H_2_O_2_ accumulation was similar between both groups. By 48 hours post-inoculation, the BO-treated plants showed a 1.5-fold increase in H_2_O_2_ accumulation compared to the CO plants. At 72 hours post-inoculation, BO-treated plants exhibited a significantly higher H_2_O_2_ accumulation of 40% of leaf area, which was nearly twice as much as the 21% seen in the CO group. These results suggest that BO-treated plants have a more robust oxidative response to the pathogen challenge.

### Comparison of Superoxide Dismutase (SOD) Accumulation in Co and Bo Plants

Histochemical analysis of SOD accumulation revealed a notable increase in SOD levels in the BO-treated plants linked to the CO control at all time points post-inoculation (0, 24, 48, and 72 hours). (Fig. 4 D) At the initial time point (0 hours), there was negligible SOD accumulation in both BO and CO samples. However, by 24 hours, BO plants exhibited significantly higher SOD levels (25%) compared to CO plants (15%). This difference became more pronounced at 48 hours, where the BO-treated samples showed 45% accumulation, compared to 30% in CO. At 72 hours, the SOD accumulation in BO reached 70%, which was notably higher than the 50% observed in CO samples. The statistical analysis confirmed that the differences between BO and CO were significant at all post-inoculation time points, with p-values of 0.0023, 0.0027, and 0.0029 at 24, 48, and 72 hours, respectively.

### Microbial Consortium-Triggered Activation of Defense and Growth-Regulating Genes in Tomato

To investigate the impact of the microbial consortium (BO) treatment on the expression of defense-associate genes in tomato, quantitative PCR analysis was performed using three target defense genes (Pto Kinase, PR2b, Chi3) and one reference gene (Ubiquitin). The expression levels of these genes were standardized to the Ubiquitin reference gene and compared between BO-treated plants and the control (CO) group.

The expression of Pto Kinase in BO-treated plants showed a modest increase of 1.2-fold compared to the control (CO) plants (Fig. 5), indicating a slight upregulation in response to the microbial consortium. In contrast, PR2b displayed a 2-fold surge in expression in BO-treated plants compared to CO, highlighting enhanced activation of defense mechanisms. The Chi3 gene showed the highest upregulation, with a 2.73-fold increase in BO-treated plants relative to CO, suggesting a robust response associated with chitinase activity, which shows a major role in plant defense.

**Figure 5:**
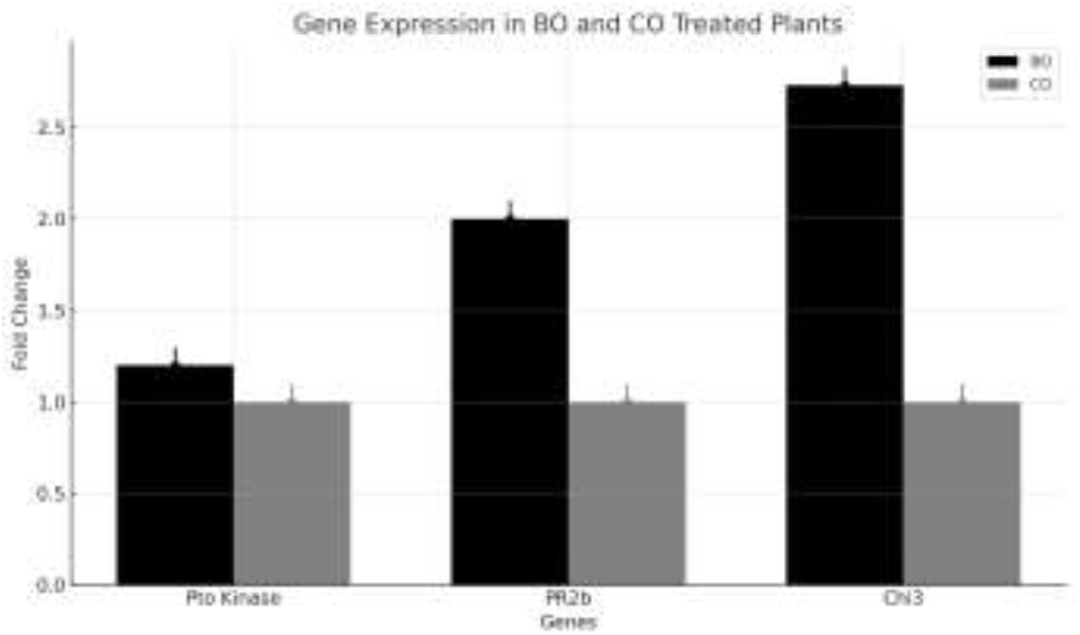
Expression of defense-related genes (Pto Kinase, PR2b, Chi3) in BO-treated and CO-control tomato plants. Data are presented as fold changes comparative to the expression in CO-control plants. Error bars signify the standard deviation of triplicate samples.

Overall, the Bo treatment led to a broad upregulation of both defense-related and growth-promoting genes, indicating its potential role in enhancing resistance to *A. solani* and promoting overall plant health.

### Comparative Metabolite Profiling of Microbial Consortium-Treated and Control Tomato Leaves

A comparative un-targeted metabolite profiling of microbial consortium-treated (BO) and control (CO) leaves was performed to elucidate the changes in metabolic level. After seven days post-treatment, leaf tissues from both groups were harvested and processed to obtain data using LC–MS/MS spectra in both positive and negative ion modes to maximize peak intensity and detection sensitivity.

The metabolic profile showed significant compositional changes between the BO and CO samples. This analysis revealed metabolites that were both unique and common to both groups, indicating a notable shift in the metabolite landscape due to the microbial consortium treatment.

Functional annotation and pathway enrichment analysis of the differentially expressed metabolites indicated significant changes in key metabolic pathways. These included pathways related to plant defense, such as phenylpropanoid biosynthesis, and growth-regulatory pathways, highlighting the impact of the microbial consortium on both defense and growth processes in the treated plants. Furthermore, statistical and chemometric analyses, including PCA and OPLS-DA, identified signature biomarker metabolites that were significantly upregulated in the Bo group, suggesting their potential involvement in developing plant resistance to *A. solani*.

### Univariate and Multivariate Data Analyses

#### Univariate analysis

Univariate analysis exposed significant metabolic alterations between the microbial consortium-treated (BO) and control (CO) tomato leaves. A t-test identified a total of 371 significant metabolite features with a p-value ≤ 0.05 (Fig. 6 A). Fold change investigation further identified 367 notably upregulated and 399 downregulated metabolite structures (Fig. 6 B). These features were distinctly visualized in a volcano plot, highlighting the differential metabolomic response between the Bo and Co samples (Fig. 6 C). This analysis underscores the metabolic shifts induced by the microbial consortium treatment, reflecting the biochemical adaptations associated with enhanced plant defense and growth.

**Figure 6:**
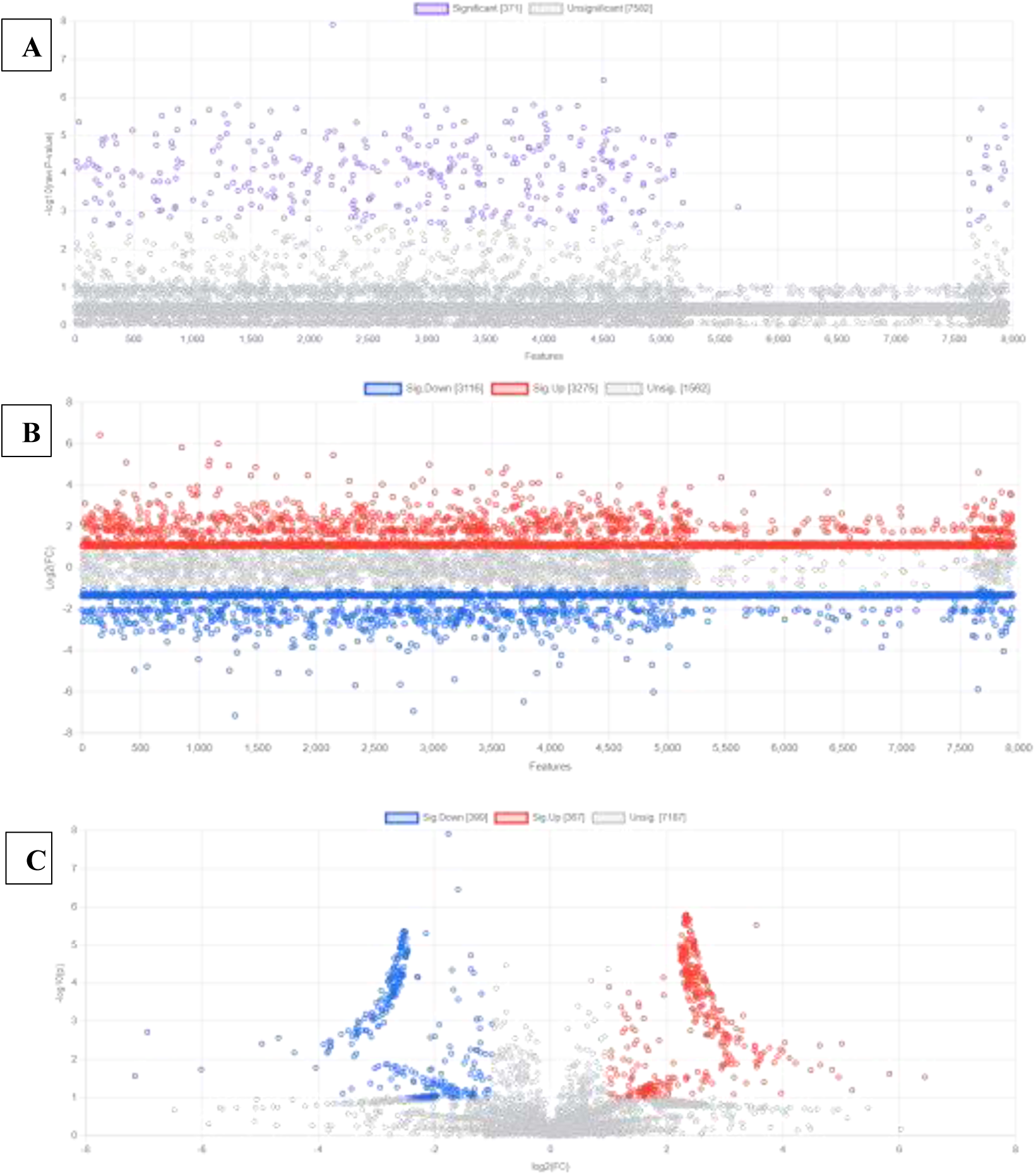
(A) Univariate analysis using a t-test identified significant metabolite features in microbial consortium-treated (BO) and control (CO) tomato leaves. A total of 371 metabolite features were significant (P ≤ 0.05), indicating substantial metabolic differences between the two groups. (B) Fold change analysis of metabolite features between BO and CO groups. Out of the identified metabolites, 367 features were significantly upregulated (red), and 399 were downregulated (blue) with a fold change threshold of ≥ 2.0, demonstrating the impact of microbial consortium treatment. (C) Volcano plot illustrating the differential expression of metabolites between BO and CO samples. Significantly upregulated (red) and downregulated (blue) metabolite features are displayed, highlighting the biochemical responses induced by the microbial consortium treatment.

### Key Metabolomic Changes in Response to Microbial Consortium Treatment

Metabolomic analysis of tomato leaves treated with the microbial consortium (BO) compared to the control (CO) revealed significant shifts in the concentration of key metabolites. The top upregulated and downregulated metabolites are presented in Table 4. Notably, metabolites such as (2E,4E)-5-(2-Furyl)-2,4-pentadienal (Fold Change = 86.94) and 2-(2,4-dihydroxyphenyl)-3,5,7-trihydroxy-4H-chromen-4-one (Fold Change = 65.09) were significantly upregulated in BO-treated samples, indicating their possible role in enhancing plant defense responses. In contrast, metabolites like 2-C-(2-Carboxyethyl)-3-deoxypentaric acid (Fold Change = 0.007) and Caffeic acid (Fold Change = 0.008) were markedly downregulated, suggesting potential suppression of certain metabolic pathways in treated plants.

**Table 4:**
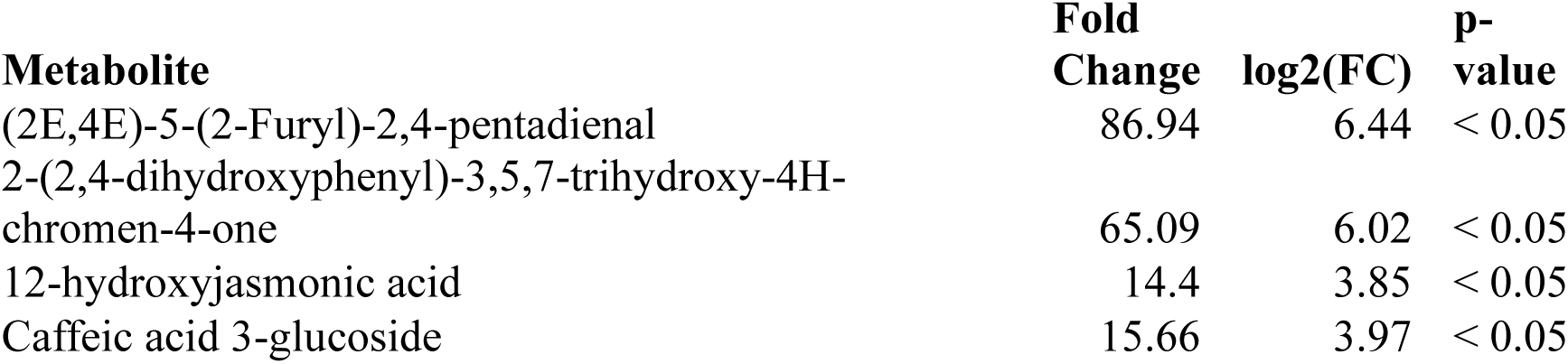
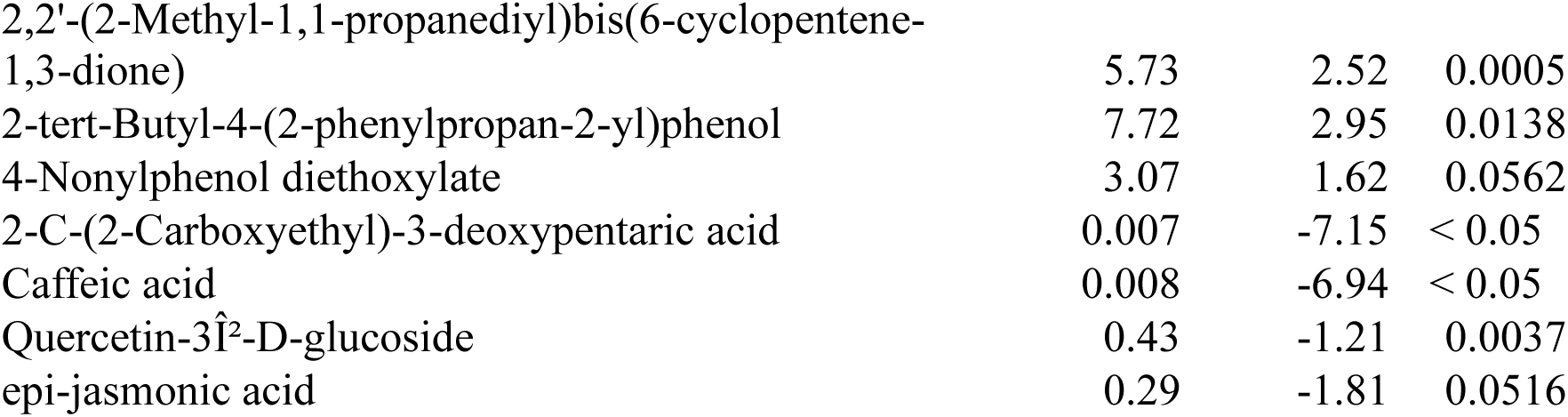
Top Upregulated and Downregulated Metabolites Based on Fold Change. Top upregulated and downregulated metabolites based on fold change, comparing microbial consortium-treated (BO) and control (CO) tomato leaves. This table highlights the metabolites with the highest positive and negative fold changes, indicating significant shifts in metabolite abundance in response to microbial treatment. The p-values reflect the statistical significance of these changes.

### Defense-Related Metabolites

Further analysis identified several metabolites specifically related to plant defense mechanism. Among these, 12-hydroxyjasmonic acid (Fold Change = 14.40) and Caffeic acid 3-glucoside (Fold Change = 15.66) were significantly upregulated in BO-treated plants. These metabolites are involved in critical defense pathways, with 12-hydroxyjasmonic acid playing a key role in the jasmonic acid signaling pathway, which is crucial for responding to pathogen attacks. Additionally, Caffeic acid 3-glucoside is known for its antioxidant properties, which contribute to the plant’s ability to mitigate oxidative stress caused by pathogen invasion.

On the other hand, defense-related metabolites such as Quercetin-3β-D-glucoside (Fold Change = 0.43) and epi-jasmonic acid (Fold Change = 0.29) were downregulated in BO-treated samples. While both are typically associated with plant defense and stress responses, their downregulation may indicate a shift in the plant’s metabolic strategy in response to the microbial consortium, potentially prioritizing other pathways for enhanced resistance.

These findings underscore the significant metabolic reprogramming induced by microbial consortium treatment, highlighting both the upregulation of key defense metabolites and the potential modulation of other metabolic pathways.

### Multivariate Analysis

#### Principal Component Analysis (PCA)

Unsupervised PCA was employed to discern patterns of metabolic variation between BO and CO samples. The PCA score plot revealed clear clustering, with variance explained by the first two principal components (PC1 and PC2) accounting for 28.5% and 20% of the total variance, respectively (Fig. 7 A). The distinct separation of the Bo and Co groups in the PCA plot indicates substantial metabolomic differences attributable to the microbial consortium treatment.

**Figure 7:**
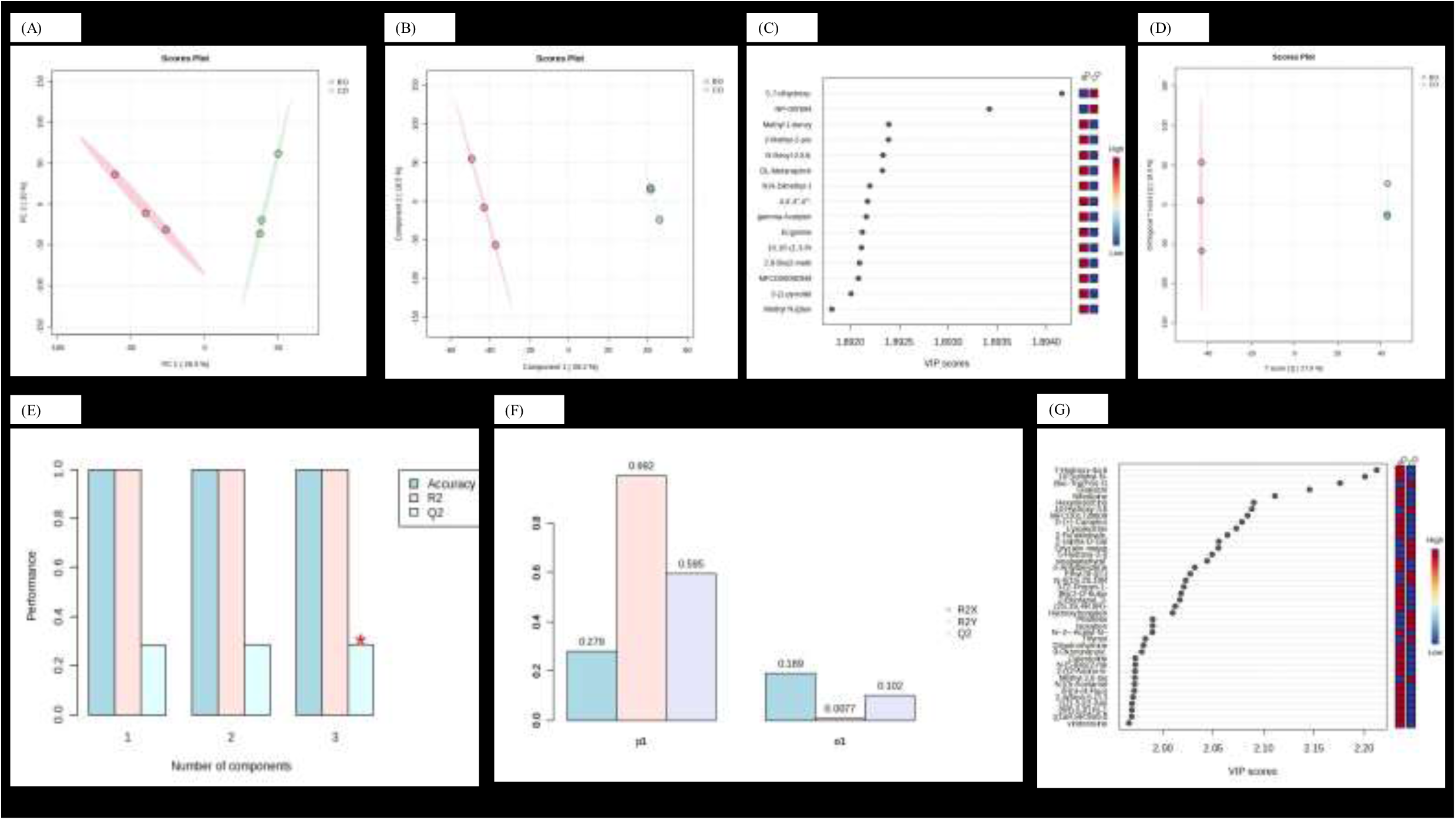
(A) Principal Component Analysis (PCA) score plot showing the distribution of BO and CO samples along the first two principal components (PC1 and PC2), demonstrating distinct separation between groups, indicative of different metabolomic profiles influenced by the microbial consortium treatment. (B) Partial Least Squares Discriminant Analysis (PLS-DA) score plot depicting discrimination between BO and CO samples. The model shows 32% covariance at component 1 (x-axis) and 17.5% covariance at component 2 (y-axis), highlighting metabolic differences between the treated and control groups. (C) Cross-validation of the PLS-DA model, with positive Q² values reflecting the models predictability and lack of overfitting, demonstrating its reliability in distinguishing BO from CO samples. (D) Variable Importance in Projection (VIP) scores for the top 25 metabolite features contributing to the PLS-DA model. Features with VIP scores ≥ 1 are highlighted as major contributors to the metabolic separation between BO and CO samples. (E) Orthogonal Partial Least Squares Discriminant Analysis (OPLS-DA) score plot illustrating separation between BO and CO groups. The T score [1] explains 27.9% of the variance, while the orthogonal T score [1] accounts for 18.9%, indicating clear metabolomic distinctions. (F) Cross-validation of the OPLS-DA model. Positive Q² values indicate strong predictability and minimal overfitting, validating the robustness of the OPLS-DA model in differentiating the metabolic profiles of BO and CO samples. (G) VIP scores of the top metabolites identified in the OPLS-DA model. High VIP scores (≥ 1) indicate key metabolites that contribute strongly to metabolic variation between BO and CO groups, potentially serving as biomarkers for microbial consortium treatment effects.

#### Partial Least Squares Discriminant Analysis (PLS-DA)

To further explore these metabolic differences, a supervised PLS-DA model was constructed, which showed significant discrimination between the Bo and Co samples with a covariance of 32% at component 1 (x-axis) and 17.5% at component 2 (y-axis) (Fig. 7 B). Cross-validation analysis of the PLS-DA model yielded positive Q² values, indicating high predictability and minimal overfitting (Fig. 7 C). The variable importance in projection (VIP) scores highlighted the top 25 metabolite features (VIP scores ≥ 1) that contributed most significantly to the observed group separation (Fig. 7 D).

#### Orthogonal Partial Least Squares Discriminant Analysis (OPLS-DA)

To enhance the interpretability of the metabolic variations, OPLS-DA was performed. The OPLS-DA score plot revealed a clear distinction between the Bo and Co groups, with a T score [1] explaining 27.9% of the variance and an orthogonal T score [1] accounting for 18.9% (Fig. 7 E). Cross-validation of the OPLS-DA model confirmed its reliability, with positive Q² values indicating high predictive accuracy (Fig. 7 F). The (Fig. 7 G) represents VIP scores for the top metabolites identified in the OPLS-DA model. Metabolites with high VIP scores (≥ 1) are key contributors to the metabolic differences between BO and CO groups and may serve as biomarkers for the effects of microbial consortium treatment.

### Biomarker Identification for Early Blight Resistance

To identify potential metabolite biomarkers associated with the microbial consortium’s impact on tomato plant resistance to early blight, a multivariate ROC curve-based exploratory investigation was accomplished. This approach allowed for the automated identification of the most relevant features. Using the PLS-DA algorithm and typical importance measures, the top 25 signature metabolite biomarkers were identified. These metabolites exhibited differential abundance in the microbial consortium-treated (Bo) versus control (Co) groups, indicating their probable role in mediating plant immune mechanisms.

The analysis revealed significant differences in the abundance of various metabolites between the microbial consortium-treated (BO) and control (CO) tomato leaves, as indicated by the Variable Importance in Projection (VIP) scores derived from the OPLS-DA model.

Among the top-ranked metabolites (Table 5), 1-Stearoyl-2-hydroxy-sn-glycero-3-PE exhibited a notable increase in the CO samples, suggesting its potential role in maintaining membrane integrity under control conditions. In contrast, several metabolites were significantly upregulated in the BO samples, indicating enhanced metabolic responses associated with the microbial consortium treatment.

**Table 5:**
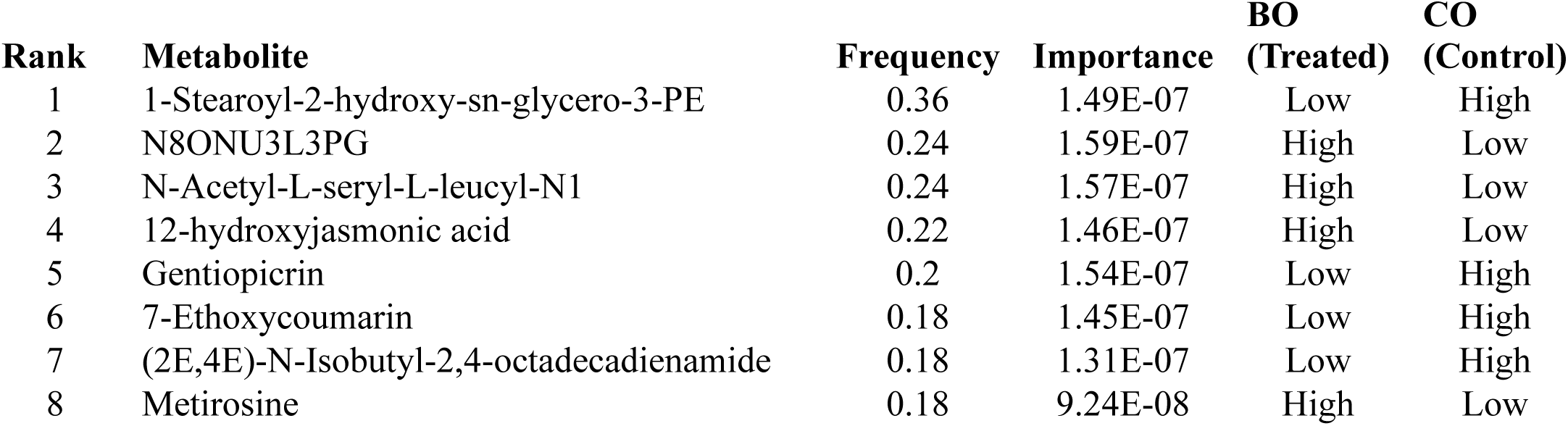
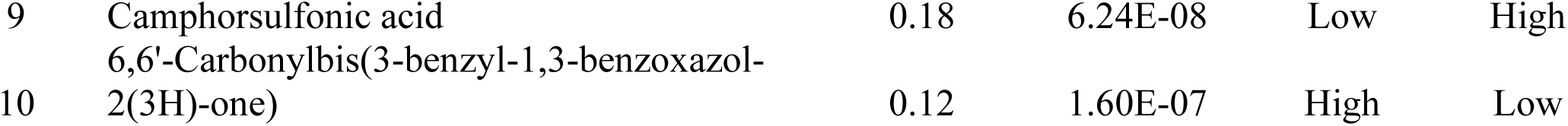
Top-ranked metabolites identified through VIP scores from the OPLS-DA model, comparing microbial consortium-treated (BO) and control (CO) tomato leaves. Metabolites with high VIP scores (≥ 1) contribute strongly to the metabolic differences between the two groups. The relative abundance of each metabolite is indicated as "High" or "Low" for BO and CO samples.

For instance, N8ONU3L3PG and N-Acetyl-L-seryl-L-leucyl-N1 were identified as key metabolites with high VIP scores (≥ 1), which may contribute to improved plant resilience and defense mechanisms. Additionally, 12-hydroxyjasmonic acid, known for its role in plant stress responses, was also found to be elevated in the BO-treated samples, further supporting the hypothesis that microbial treatments enhance stress resilience in tomato plants.

Furthermore, metabolites such as Metirosine and 3-Hydroxybufa-14,20,22-trienolide showed significantly lower levels in the CO samples, highlighting their potential as biomarkers for the effects of microbial consortium treatment. The distinct metabolic profiles observed between BO and CO samples underscore the influence of microbial treatments on tomato leaf physiology, suggesting that these metabolites could serve as indicators of the metabolic adaptations facilitated by the microbial consortium.

Overall, the differential abundance of these metabolites illustrates the profound impact of the microbial consortium on the metabolic pathways in tomato plants, paving the way for future investigations into their functional roles and potential applications in enhancing plant health and productivity.

#### Biomarker Annotation

The identified biomarker metabolites were further annotated using databases such as MassBank and the Plant Metabolic Network (PMN) specific to *Solanum lycopersicum*. The annotation process led to the identification of several key metabolites. Among these, metabolites such as N8ONU3L3PG, 3-Methyladenine, 12-hydroxyjasmonic acid, Trifolin, and Isorhamnetin exhibited significant changes in abundance between BO and CO samples. Notably, some of these biomarkers, including 3-Methyladenine and 12-hydroxyjasmonic acid, were upregulated in Bo, indicating a possible enhancement of defense-related metabolic pathways in the microbial consortium-treated plants.

#### Class Probabilities and Metabolite Importance

The ROC curve analysis also included a probability view to show the predicted class probabilities for Bo and Co samples, highlighting clear distinctions in their metabolomic profiles (Fig. 8B). The metabolites with the highest importance, such as 1-Stearoyl-2-hydroxy-sn-glycero-3-PE, Gentiopicrin, and Camphorsulfonic acid, contributed significantly to the separation between Bo and Co groups (Fig. 8A). These metabolites are likely key players in the biochemical pathways associated with improved resistance to early blight in tomato plants. Among the biomarker metabolites, the up-regulated and downregulated during the treatment has been plotted in (Fig. 8C-H).

**Figure 8:**
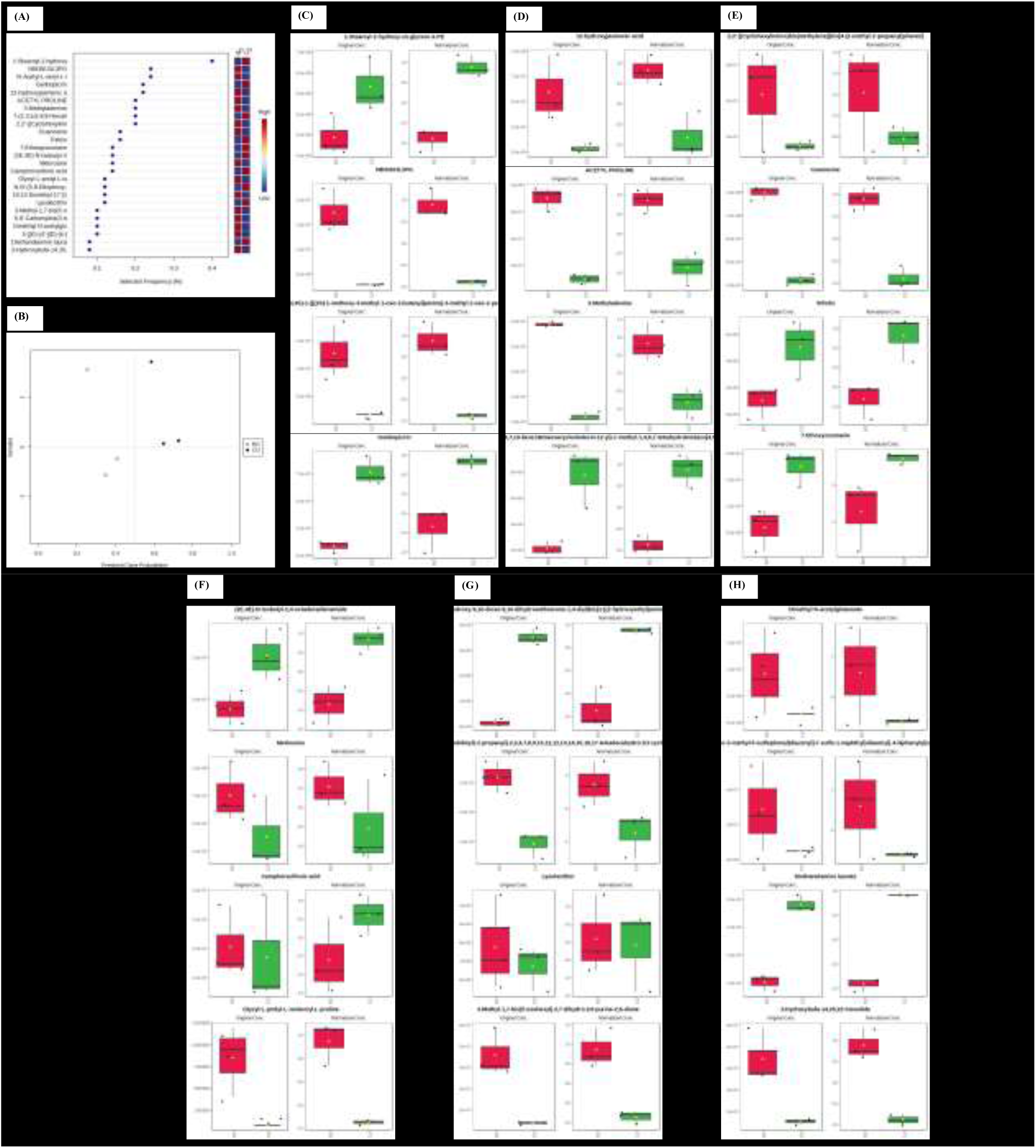
(A) Multivariate exploratory receiver operating characteristic (ROC) curve analysis showcasing the top 25 potential metabolite biomarkers identified through the Partial Least Squares Discriminant Analysis (PLS-DA) algorithm. The metabolites highlighted in this analysis significantly contribute to the differentiation between the microbial consortium-treated (Bo) and control (Co) groups. (C) Predicted class probabilities derived from the ROC curve analysis, demonstrating clear metabolomic distinctions between the Bo and Co samples. Open circles (●) denote Bo samples, while closed circles (●) indicate Co samples. Annotated down and upregulated biomarkers are illustrated in 8(C-H).

### Analysis of Metabolic Pathway Enrichment and Associated Impact

Annotated metabolites with KEGG identifiers were exposed to KEGG Mapper for pathway mapping, revealing their improvement across various metabolic pathways. The analysis identified a total of 50 enriched metabolic pathways in the *S. lycopersicum* (sly) database. The most prominent pathways included "metabolic pathways" and "biosynthesis of secondary metabolites," each showing significant hits, indicating the diverse metabolic changes triggered in the treated plants (Figure 9 A). Other notable pathways with significant hits were related to tryptophan metabolism, tyrosine metabolism, riboflavin metabolism, purine metabolism, and amino acid biosynthesis.

**Figure 9:**
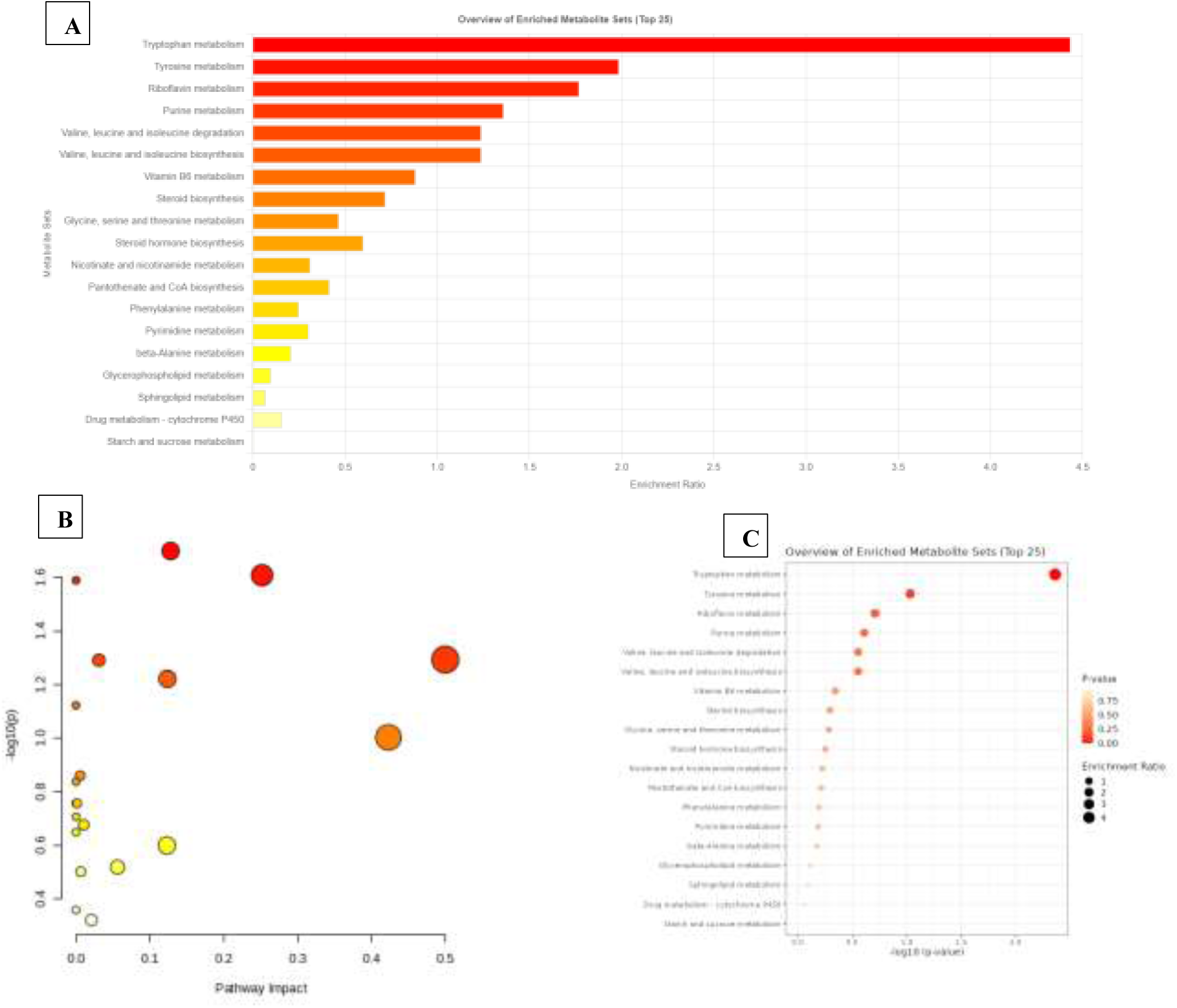
(A) Pathway mapping illustrating the enriched metabolic pathways in microbial consortium-treated tomato plants. The length of each bar corresponds to the number of metabolite hits identified within each pathway, highlighting the pathways most affected by treatment. (B) Pathway impact analysis assessing the significance of each pathway in the tomato plant’s response to treatment. The size of the circles reflects the pathway impact, while the color gradient, ranging from red to yellow, indicates the level of statistical significance associated with each pathway. (C) Distribution of metabolite features obtained from a fold change (FC) threshold of ≥ 2.0 categorized into different compound groups using MetaboAnalyst 6.0. The circles represent various chemical classes, with circle size denoting the enrichment ratio and color indicating the p-value, demonstrating the diversity and significance of annotated metabolites across chemical classifications.

Pathway impact (PI) analysis evaluated the influence of these pathways on plant responses. The analysis integrates centrality and enrichment results to highlight pathways with major impact. Significant pathways (p ≤ 0.05) showing major impact in tomato plants included tryptophan metabolism, tyrosine metabolism, riboflavin metabolism, purine metabolism, and amino acid biosynthesis, indicating their crucial roles in plant defense and metabolic regulation under the treatment (Figure 9 B).

In the pathway analysis, distribution of the identified metabolites across different chemical classes was conducted. The enrichment analysis grouped these metabolites into key chemical classes such as monosaccharides, isoprenoids, sterols, peptides, fatty acids, prenol lipids, amino acids, and conjugates all displaying statistically notable p-values (P ≤ 0.05) (Figure 9 C). Among these, isoprenoids, fatty acids, sterols, and amino acids were prominently represented, suggesting their prominent roles in plant tolerance mechanisms and metabolic adjustments under the treatment conditions.

Metabolomic analysis further suggested some overlaps in the metabolite datasets, reflecting the complex and interconnected nature of plant metabolism. This overlap adds depth to the metabolomic data, making it more representative and insightful for understanding the metabolic changes in treated tomato plants.

This enrichment and impact analysis provide a comprehensive overview of the metabolic alterations in tomato plants, highlighting the involvement of key metabolic pathways and

### Heatmap for Metabolite Profiling

Hierarchical clustering analysis revealed distinct metabolic profiles between BO-treated and CO-treated tomato leaves. The heatmap shows clear separation between the two sample groups, BO (treated) and CO (control), based on the relative abundance of metabolites (Figure 10). The clustering dendrogram on the top reflects the separation of the BO and CO groups, highlighting that BO treatment induced significant metabolic reprogramming. BO-treated samples (BO1, BO2, BO3) clustered together, showing a different metabolic pattern compared to the CO group (CO1, CO2, CO3).

**Figure. 10.**
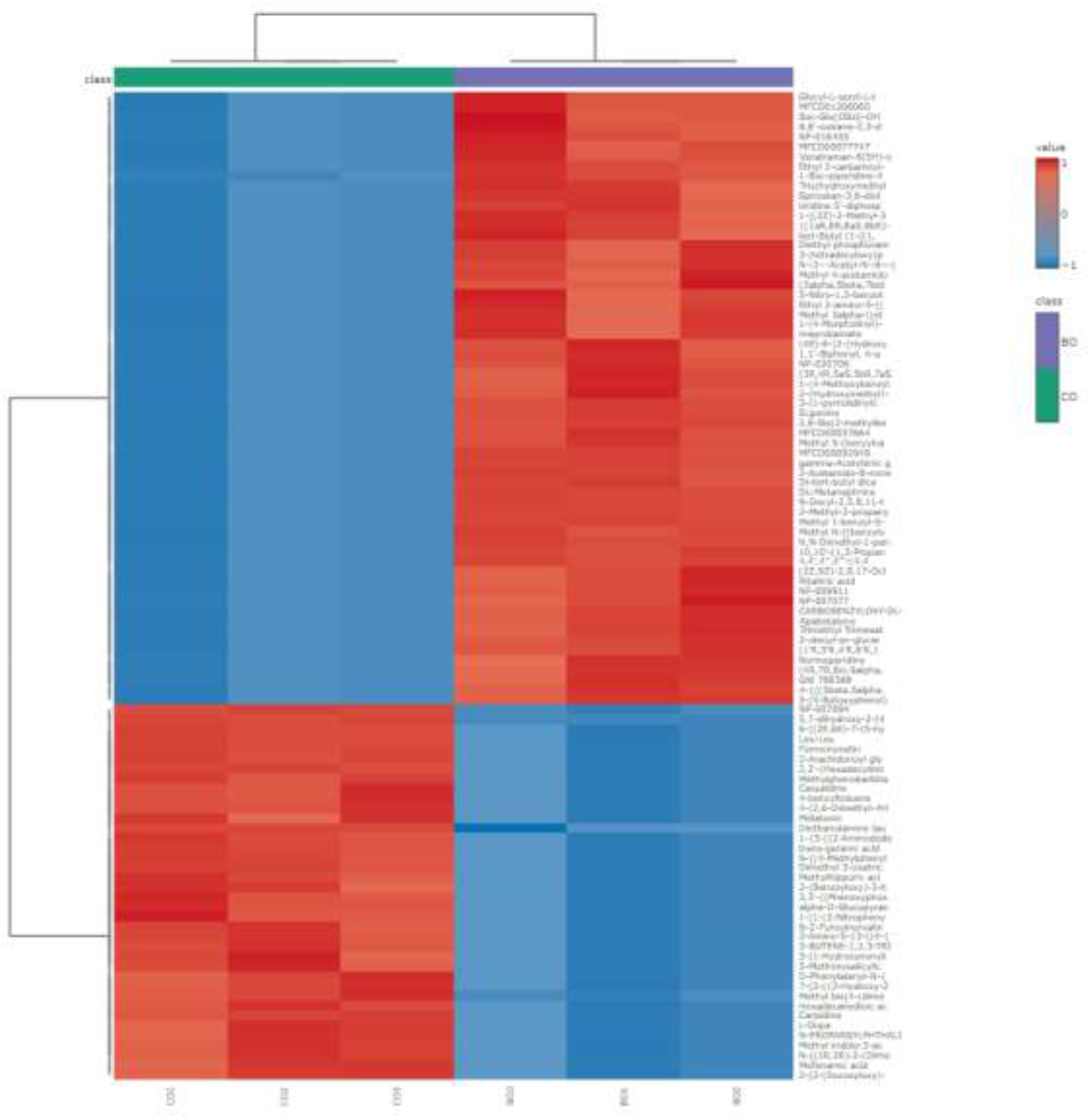
Hierarchical clustering was performed on metabolite feature abundances, which were normalized through Pareto scaling and subjected to a t-test from the adjusted dataset. Pearson correlation was utilized as the distance metric, and Ward’s method was applied for the clustering process. The heatmap shows the relative richness of the top 50 metabolite features in BO-treated (BO1, BO2, BO3) and CO-treated (CO1, CO2, CO3) samples. The color scale represents the relative abundance, from higher (red) to lower (blue). Dendrograms at the top show the hierarchical relationships between BO and CO groups, while the dendrogram on the left displays the clustering of identified metabolite features.

Metabolites that were more abundant in BO-treated plants are represented by intense red in the heatmap, while metabolites with lower abundance in BO (and higher in CO) are indicated by blue. Key metabolites, such as Glycyl-L-seryl-L, Trifolin, and Camphor showed higher abundance in BO-treated samples, indicating a strong response to microbial consortium treatment. Conversely, metabolites like Uridine and Apabetalone displayed lower levels in BO compared to CO.

These results suggest that the microbial consortium treatment (BO) triggered specific metabolic shifts that differentiate BO-treated plants from the control group, implying a broad metabolic reprogramming in response to the treatment.

## Discussion

### Nutrient Composition of Gir Cow Dung and Its Agricultural Implications

The nutrient analysis of Gir cow dung manure reveals a highly favorable profile for agricultural use. Cow dung’s neutral pH, adequate moisture content, and high nutrient levels (including nitrogen, phosphate, and potash) make it effective as a natural fertilizer and soil conditioner (Kadam et al. 2024). Studies confirm that cow dung enriches soil with essential nutrients and enhances microbial activity, thus supporting sustainable agriculture (Gupta, Aneja, and Rana 2016). Furthermore, its use as an organic amendment to biomass has been shown to improve soil physio-chemical properties, like pH, electrical conductivity, and the carbon-to-nitrogen ratio, making it a promising organic solution for increasing soil fertility (Bassem Yassin et al. 2021).

### Bacterial Diversity and Its Role in Enhancing Plant Disease Resistance

The 16S rRNA amplicon sequencing of Gir cow dung revealed a rich microbial community, predominantly composed of beneficial genera such as *Lactococcus*, *Bacillus*, *Clostridium*, and *Sorangium*. Genus level, *Lactococcus* was the prominent (43.9%), followed by *Clostridium* and *Bacillus*, with *Bacillus* species known for their antifungal properties and plant growth-promoting effects. Recent research has confirmed the versatile role of *Bacillus* spp. in promoting plant health through several mechanisms, including solubilization of phosphate, release of phytohormones like indole-3-acetic acid (IAA), and siderophore secretion, which enhance nutrient uptake and stress tolerance (Vasques, Nogueira, and Hungria 2024).

The *Bacillus* species isolated in this study, including *Bacillus licheniformis*, *Bacillus subtilis*, and *Bacillus tequilensis*, demonstrated significant antagonistic activity against *A. solani*, aligning with recent findings where *Bacillus* strains showed strong biocontrol capabilities by producing antifungal metabolites like iturin and fengycin (Vasques, Nogueira, and Hungria 2024; W. Chen et al. 2023). These metabolites disrupt the cell wall of pathogenic fungi, contributing significantly to plant resistance.

*Lactococcus* spp., which dominated the microbial community, play a prominent role in enhancing plant health through nutrient cycling and plant hormone production. *Lactococcus lactis*, for instance, has been reported to facilitate biological nitrogen fixation and solubilize phosphorus, enhancing nutrient availability to plants (Jaffar, Jawan, and Chong 2023). Moreover, lactic acid bacteria (LAB) like *Lactococcus* contribute to pathogen suppression by producing antimicrobial compounds, creating an unfavorable environment for pathogens and bolstering plant defenses (Lamont et al. 2017).

The combination of *Lactococcus*, *Bacillus*, and other beneficial microbes in Gir cow dung underscores its potential as a natural bioresource for improving plant health through both growth promotion and pathogen suppression. The synergistic interactions between these microbial species enhance plant resilience, reduce dependence on chemical pesticides, and contribute to sustainable agricultural practices (Pellegrini et al. 2024)

### Biocontrol Potential of *Bacillus* Isolates

The *Bacillus* isolates tested, including *B. subtilis*, *B. licheniformis*, and *B. tequilensis*, exhibited significant antagonistic activity against *Alternaria solani*. This inhibition aligns with prior studies that demonstrated the antifungal properties of *Bacillus* species, which are often attributed to their release of secondary metabolic compounds, like lipopeptides and phenolic products (Bonaterra et al. 2022; Gao et al. 2021). *Bacillus subtilis* ZD01 has been previously reported to exhibit strong antagonistic properties against *A. solani* (D. Zhang et al. 2020). *B. licheniformis* NCIMB 8874 produces the antifungal peptide ComX, which is effective against the fungal leaf pathogen *A. alternata* (Shleeva, Kondratieva, and Kaprelyants 2023). However, no reports have been found so far indicating its antifungal activity against *A. solani*. *B. tequilensis* has been shown to share similar biosynthetic gene clusters and secondary metabolites with *B. subtilis* ZD01 (Karun and Arunachalam 2023), there are currently no reports of its antagonistic activity against *A. solani*. A study by (Zahoor et al. 2022) reports that *B. tequilensis* reduced disease severity in tomato plants infected by *A. alternata*. The efficacy of these isolates as biocontrol suppressors underscores their suitability for use in sustainable agricultural practices to manage plant diseases and develop plant health.

### Plant Growth-Promoting Traits and Biochemical Characterization

The biochemical depiction of the *Bacillus* species showed plant growth-enhancing abilities like siderophore release, nitrogen fixation, phosphate solubilization, and IAA production, which are crucial for nutrient availability and plant vigor. These traits are supported by findings from (Shi et al. 2024), who reported that microbial inoculants significantly improved crop development over mechanisms like phosphate solubilization and IAA production (Shi et al. 2024). *Streptomyces sp. NEAU-S7GS2*, another microbial agent, also displayed phosphate solubilization, IAA production, and antagonistic activity against *Sclerotinia sclerotiorum*, further highlighting the potential of beneficial microbes in promoting plant devolopment and health (D. Liu et al. 2019).

### Enhanced Disease Resistance in BO-Treated Plants

BO-treated plants exhibited significantly lower lesion formation in response to *A. solani* infection, and higher levels of hydrogen peroxide (H₂O₂), which suggests an enhanced oxidative burst a key defense response. Similar effects have been reported by (W. Chen et al. 2023) where *Bacillus subtilis* was found to induce an oxidative response in treated plants, boosting their resistance to pathogens (W. Chen et al. 2023). The initiation of systemic acquired resistance (SAR) and other defense mechanisms is consistent with reports indicating that *Bacillus* species, along with other microbial inoculants, can effectively prime plant defenses, making them highly resistant to subsequent pathogen attacks (Bonaterra et al. 2022)

### Expression of Defense and Growth-Related Genes

The upregulation of genes related to defense, like *Pto kinase*, *PR2b*, and *Chi3*, in BO-treated plants further confirms the role of microbial consortia in enhancing plant immunity. The significant upregulation of the *Chi3* gene, associated with chitinase production, is consistent with observations by (Gao et al. 2021), where *Bacillus velezensis* strains were shown to produce chitinase, a key enzyme for fungal cell wall degradation (Gao et al. 2021). Additionally, *Streptomyces* species have also demonstrated the capacity to enhance plant immunity by secreting enzymes like glucanase and cellulase, which contribute to pathogen suppression (D. Liu et al. 2019).

### Metabolomic Alterations in Tomato Plants Treated with Microbial Consortium

The metabolomic profiling of microbial consortium-treated (BO) and control (CO) tomato leaves revealed significant metabolic reprogramming. The distinct metabolomic profiles, detected via LC-MS/MS together with +ve and -ve ion modes, highlight the impact of the microbial consortium treatment in altering key metabolites involved in plant defense as reported by (Vishwakarma et al. 2020). These changes underscore the role of microbial interactions in enhancing resistance against phytopathogens by modulating various metabolic pathways in tomato leaves.

### Differential Metabolic Features and Functional nrichment Analysis

In this study, univariate analysis identified 371 significantly altered metabolite features (p ≤ 0.05) between the Bo and Co groups. The Volcano Plot analysis revealed that 367 metabolites were significantly upregulated, while 399 metabolites were significantly downregulated in response to the microbial consortium treatment. This significant differential expression points to a broad shift in the metabolic landscape, possibly reflecting the plant’s adaptive responses to biotic stress. For instance, research has shown that plants under biotic stress undergo significant shifts in metabolism, including the accumulation of defense-related compounds like phenolics, terpenoids, and phytohormones (Anzano et al. 2021). The notable increase in metabolites such as (2E,4E)-5-(2-Furyl)-2,4-pentadienal (Fold Change = 86.94) and 2-(2,4-dihydroxyphenyl)-3,5,7-trihydroxy-4H-chromen-4-one (Fold Change = 65.09) suggests their potential involvement in key metabolic processes within the plant. Although their precise role in defense mechanisms remains unclear, their significant upregulation in response to microbial consortium treatment indicates that these compounds may contribute to broader metabolic reprogramming. Additionally, 12-hydroxyjasmonic acid (Fold Change = 14.40) and Caffeic acid 3-glucoside (Fold Change = 15.66) were found to be significantly upregulated, indicating the activation of jasmonic acid signaling and antioxidant pathways, both of which are crucial for the plant’s response to pathogen attack (Yang et al. 2019; Monteiro Espíndola et al. 2019).

On the other hand, metabolites such as 2-C-(2-Carboxyethyl)-3-deoxypentaric acid (Fold Change = 0.007) and caffeic acid (Fold Change = 0.008) were markedly downregulated, which may suggest the suppression of certain metabolic pathways, potentially reallocating resources towards more critical defense mechanisms in Bo-treated plants. The downregulation of quercetin-3β-D-glucoside (Fold Change = 0.43) and defense-related metabolites like epi-jasmonic acid (Fold Change = 0.29) might indicate a shift in the plant’s metabolic strategy, prioritizing other metabolic pathways that enhance resistance through alternative mechanisms. (+)-7-iso-Jasmonoyl-L-isoleucine (epi-jasmonic acid) serves as a bioactive form of jasmonates in plant defense, effectively activating jasmonate signaling pathways by binding to the COI1 receptor. While epi-jasmonic acid is capable of initiating defense responses, its activity is moderate compared to other jasmonate derivatives. Its stereoisomeric structure allows for effective receptor interaction, but it may not fully optimize defense responses under all stress conditions, indicating that other jasmonate forms like 12-hydroxyjasmonic acid may play more prominent roles in certain defense mechanisms (Fonseca et al. 2009).

### Multivariate Analyses

Multivariate analyses, including Principal Component Analysis (PCA), Partial Least Squares Discriminant Analysis (PLS-DA), and Orthogonal PLS-DA (OPLS-DA), provided strong evidence of the distinct metabolic profiles between Bo-treated and control (Co) tomato leaves (Kim et al. 2017). PCA revealed significant variance, with the first two components (PC1 and PC2) explaining 28.5% and 20% of the variance, respectively. This supports the hypothesis of widespread metabolic reprogramming induced by the microbial consortium treatment.

The positive Q² values indicated that these PLS-DA and OPLS-DA models are robust and not prone to overfitting, making them reliable for distinguishing the metabolic shifts between treatments. Variable Importance in Projection (VIP) scores identified the top 25 metabolites contributing most significantly to group separation, reinforcing the notion that specific metabolic changes are key drivers of the observed differences (Wang et al. 2020). However, at this stage, the focus remains on the statistical validation of metabolic divergence between the groups, without delving into the functional implications of these metabolites.

### Key Metabolic Pathways Impacted by Microbial Treatment

The analysis of KEGG pathway enrichment and functional annotation for the differentially expressed metabolites pointed to significant changes in key metabolic pathways related to both plant defense and growth regulation. Notably, pathways such as biosynthesis of phenylpropanoid, flavonoid, and terpenoid backbone were significantly enriched in Bo-treated plants, indicating their crucial roles in bolstering plant defense mechanisms (Vogt 2010; Rahim, Zhang, and Busatto 2023; Xie et al. 2012). These findings align with previous studies where similar pathways were implicated in reinforcing plant structural defenses and synthesizing antimicrobial compounds in answer to biotic stress (Caulier et al. 2018).

Enrichment of phenylpropanoid biosynthesis is particularly noteworthy, as this pathway is integral to the production of lignin, flavonoids, and other phenolic compounds that contribute to cell wall reinforcement and antimicrobial defense (Yao et al. 2021). The observed upregulation of flavonoid biosynthesis suggests an increase in secondary metabolic compounds with antioxidant resources, which may show a role in reducing oxidative damage caused by *A. solani* infection (S. Chen et al. 2023). Flavonoids, such as trifolin and isorhamnetin, were prominently represented among the upregulated metabolites, supporting their role as key players in the plant’s defensive response.

### Biomarker Identification and Potential Role in Defense

Building on the multivariate analyses, a further in-depth exploration of the top metabolites identified through VIP scores was conducted to assess their biological significance in plant defense. Among the key metabolites identified were 12-hydroxyjasmonic acid, N8ONU3L3PG, and N-Acetyl-L-seryl-L-leucyl-N1, all of which were significantly upregulated in Bo-treated samples. 12-hydroxyjasmonic acid, in particular, plays a crucial role in the jasmonic acid signaling pathway, which regulates defense responses against necrotrophic pathogens such as *Alternaria solani* (Carvalhais et al. 2013).

Additionally, metabolites like Metirosine and 3-Hydroxybufa-14,20,22-trienolide were found to be significantly downregulated in the control (CO) samples, suggesting their potential as biomarkers for the microbial consortium treatment’s effects on metabolic reprogramming. The identification of these metabolites as key indicators through ROC analysis, which yielded an AUC of 1, underscores their predictive power and functional relevance in enhancing plant resilience.

Together, the differential abundance of these metabolites demonstrates the profound impact of microbial consortium treatment on tomato plant, particularly in the context of boosting the plant’s stress responses and immune functions.

### Hierarchical Clustering and Metabolic Reprogramming

The heatmap generated from hierarchical clustering analysis revealed distinct metabolic profiles between Bo-treated and Co-treated tomato leaves, highlighting the significant metabolic reprogramming induced by microbial treatment (Li et al. 2022). The clustering pattern demonstrated that Bo-treated samples grouped closely together, distinct from the Co group, underscoring the broad metabolic shifts triggered by the microbial consortium.

Key metabolites such as Glycyl-L-seryl-L, trifolin, and camphor showed higher abundance in Bo-treated plants, suggesting their roles in reinforcing structural defenses and modulating stress responses. Conversely, metabolites like uridine and apabetalone were downregulated in the Bo-treated group, which could reflect a shift in metabolic priorities from nucleic acid synthesis to defense-related processes. This shift from primary to secondary metabolism is indicative of a strategic resource allocation by the plant to support enhanced defense, a phenomenon that has been observed in other plant-pathogen interactions (Erb and Kliebenstein 2020).

### Pathway Impact and Enrichment Analysis

Pathway impact analysis provided additional insights into the importance of specific metabolic pathways in the Bo-treated plants. High-impact pathways included tryptophan metabolism, tyrosine metabolism, riboflavin metabolism, and purine metabolism, each playing crucial roles in plant defense regulation. For example, the enrichment of tryptophan metabolism indicates its involvement in synthesizing secondary metabolites like indole derivatives, which have known roles in plant immune responses (Ishihara et al. 2011; Munir et al. 2019; F. Liu et al. 2010; Dark et al. 2011).

The distribution of identified metabolites across different chemical classes, including isoprenoids, fatty acids, sterols, and amino acids, further supports the comprehensive nature of the metabolic reprogramming observed. The prominence of isoprenoids and fatty acids among the enriched metabolites suggests their role in modulating membrane dynamics and signaling processes (Jordan, Nee, and Lane 2019) that are critical for mounting an effective defense against *A. solani*.

### Implications and Future Prospects

The comprehensive metabolomic shifts observed in Bo-treated tomato plants underscore the effectiveness of the microbial consortium in inducing both defensive and growth-regulatory metabolic pathways. The identification of key biomarkers, such as 3-Methyladenine and 12-hydroxyjasmonic acid, and the significant enrichment of defense-related pathways spotlight the ability of this microbial consortium as a sustainable solution for enhancing crop resilience.

Future research should focus on the functional validation of these biomarkers to better understand their roles in the underlying mechanisms of disease resistance. Additionally, the impact of these microbial treatments in diverse environmental conditions should be evaluated to ensure their broader applicability in agricultural practices. The integration of these findings into breeding programs or foliar treatments could provide a strategic approach to drive the resistance in tomato and other crops against early blight and similar pathogens.

## Methodology

### Selection of Microbial Source

#### Collection and Processing of Gir Cow Dung

Fresh Gir cow dung samples were collected from Sri Narayani Peedam, Sripuram, Vellore, Tamil Nadu, India. The samples were collected in clean, sterile containers to prevent contamination. They were then spread under shade conditions and allowed to dry to an optimum dryness level suitable for agricultural application. The dried cow dung was homogenized to ensure consistency before further processing for nutrient analysis and microbial isolation.

#### Nutrient Analysis

The nutrient composition of the cow dung samples was analyzed at the National Agro Foundation Laboratory Services Division. The samples were sent for a comprehensive nutrient analysis to determine their suitability for microbial isolation and agricultural application. The following standardized protocols were employed: pH was assessed using a digital pH-meter; electrical conductivity was defined using a conductivity meter; moisture content was assessed by drying the sample at 105°C to constant weight. Total nitrogen was quantified using the Kjeldahl method, and total phosphate as P_2_O_5_ was measured by spectrophotometric analysis after acid digestion. Total potash as K_2_O was determined using flame photometry. Trace elements such as zinc, copper, and nickel were measured using atomic absorption spectroscopy (AAS). Total bio-carbon was analyzed with the Walkley-Black technique. Additionally, calcium, magnesium, and sodium were quantified using AAS. Particle size distribution was defined by the sieving technique, and bulk density was quantified by the core method. The color and odor of the samples were assessed visually and by sensory evaluation, respectively.

#### Metagenomic DNA Extraction, 16S rrna Amplification, and Sequencing Workflow

Metagenomic analysis was conducted on cow dung samples through a series of steps, starting with DNA extraction. This was achieved using the QIAGEN soil kit, adhering to the manufacturer’s instructions. The concentration and purity of the extracted DNA were verified using a NanoDrop spectrophotometer, aiming for 260/280 nm absorbance ratios between 1.8 and 2 for optimal results.

For the 16S rRNA bacterial region amplification, primers V13F (5’AGAGTTTGATGMTGGCTCAG3’) and V13R (5’TTACCGCGGCMGCSGGCAC3’) were utilized. PCR was performed with a reaction mixture containing 40 ng of extracted DNA, 10 pM of each primer, and Xploregen Discoveries Taq Master Mix. This master mix comprised high-fidelity DNA polymerase, 0.5 mM dNTPs, 3.2 mM MgCl2, and the necessary PCR enzyme buffer. The PCR protocol included an initial denaturation at 95°C, followed by 25 cycles of 95°C for 15 seconds, annealing at 60°C for 15 seconds, and extension at 72°C for 2 minutes, concluding with a final extension at 72°C for 10 minutes before holding at 4°C.

Post-amplification, the products were purified and assessed using a 2% agarose gel and Nanodrop quality control. AMPure beads were employed to remove any remaining primers. Sequencing libraries were constructed with an additional 8 cycles of PCR using Illumina barcoded adapters. The libraries were then purified with AMPure beads and quantified with the Qubit dsDNA High Sensitivity Assay Kit. The sequencing was carried out on an Illumina MiSeq platform using a 2x300 paired-end (PE) v3 sequencing kit.

#### Processing of Sequence Data

Raw sequencing data underwent quality control using FASTQC (v0.11.2) and MULTIQC (v1.9). Low-quality reads and adapter sequences were trimmed with TRIMGALORE. The subsequent steps involved merging paired-end reads, eliminating chimeras, and determining operational taxonomic unit (OTU) abundance. Representative sequences were selected, corrected, and classified taxonomically through the Biokart Pipeline, facilitating precise analysis at the genus level. Taxonomic assignments were made using the SILVA, GREENGENES, and NCBI databases, based on identity and coverage thresholds. The raw sequencing data are available in the NCBI’s Sequence Read Archive (SRA) database under BioSample accession SAMN44368148.

#### Statistical Evaluation

All statistical evaluations were conducted using the R statistical software (v.4.0.4, The R Foundation for Statistical Computing, Vienna, Austria). Bacterial diversity within the Gir cow dung samples was analyzed using a variety of statistical approaches. The relative abundance of the microbial community and individual taxa was computed using Microsoft Excel 2010. The formula used for calculating the relative abundance of each taxon is:

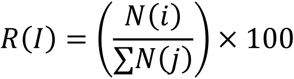

where sum∑ represents the total count across all species (j = 1 ton), N(i) is the count of the i^th^ species,

∑N(J) is the total count across all species, and

R(I) is the relative abundance of the i^th^ species as a percentage.

The analysis aimed to identify bacterial strains with potential antagonistic properties against *Alternaria solani* and those capable of enhancing plant immune responses.

#### Isolation and Screening of Microbes

Cow dung samples were serially diluted and spread onto nutrient agar plates, followed by incubation at 30°C for 48 hours. Bacterial colonies with different morphological features were selected and further purified through repeated streaking on new agar plates. This process resulted in the isolation of 50 distinct bacterial strains.

#### Assessment of Antifungal Activity Against *Alternaria solani*

To evaluate the antagonistic properties of the isolated bacteria against *A. solani*, a dual culture assay was conducted. A plug of *A. solani* mycelium was placed at the center of a potato dextrose agar (PDA) plate, while each bacterial isolate was streaked approximately 2 cm away from the fungal plug. The plates were then incubated at 28°C for 5-7 days. The degree of fungal growth inhibition was recorded and quantified. The formula used for calculating the percentage of inhibition is as follows:

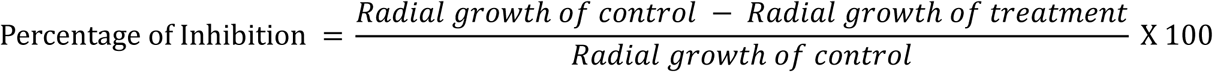

Bacterial isolates that demonstrated significant antagonistic effects, with inhibition rates exceeding 50%, were chosen for further analysis through 16S rRNA gene sequencing. This identification process was carried out at Biokart India Pvt. Ltd.

Genomic DNA extraction followed the phenol-chloroform method, and the ∼2 kb 16S rRNA gene fragment was amplified using high-fidelity PCR polymerase along with specific primers. Sequencing of the PCR products was done in both directions using the ABI 3130xl sequencing platform. The sequence data were aligned and analyzed to determine the identity of the bacterial isolates by comparing them with known sequences using BLAST and constructing a phylogenetic tree. The identified bacteria included *Bacillus licheniformis, Bacillus subtilis*, and *Bacillus tequilensis*. These three isolates, showing notable antagonistic potential, were selected for additional investigation.

The interactions between the selected *Bacillus* isolates and *A. solani* were examined using scanning electron microscopy (SEM). SEM analysis was conducted using a Carl Zeiss EVO 18 Research model. To prepare for this, co-cultures of antagonistic bacteria and *A. solani* hyphae were grown in potato dextrose broth and incubated at 28°C for 5 days. Following incubation, the hyphal structures were analyzed to visualize the physical interactions between the bacteria and the fungal pathogen. Additionally, *Alternaria* spores were cultured separately and exposed to the bacterial consortium. These interactions were also incubated at 28°C and analyzed using SEM. This detailed study allowed for a closer look at the morphological changes and interactions between the bacterial isolates and fungal structures.

#### Characterization of Antagonistic Bacteria

The three selected *Bacillus* isolates were subjected to biochemical characterization using the KB001 kit from HiMedia Laboratories. This kit was instrumental in determining a range of metabolic and physiological traits of the isolates. Tests conducted included those for indole production, methyl red reaction, citrate utilization, Voges-Proskauer reaction, and the ability to ferment various carbohydrates, such as sorbitol, glucose, lactose, arabinose, adonitol, sucrose, mannitol, and rhamnose. The kit included reagents like reagent of Kovac’s for testing the production of indole, methyl red as an indicator, and reagents of Barritt’s A and B for the Voges-Proskauer reaction. These tests allowed for a detailed understanding of the isolates’ metabolic profiles, facilitating their identification.

#### Evaluation of Plant Growth-Promoting Traits

The *Bacillus* isolate’s ability to promote plant growth was investigated through several specific assays. To assess siderophore production, the isolates were inoculated on chrome azurol S (CAS) agar plates, which were then incubated at 28°C for 48 hours. Siderophore production was indicated by the formation of an orange halo surrounding the colonies, and the size of these halos was measured to quantify production. Phosphate solubilization capacity was evaluated using Pikovskaya’s agar by placing bacterial cultures on the agar plates, followed by a 7-day incubation at 28°C. The presence of clear zones around the colonies suggested phosphate solubilization and the diameter of these zones was recorded to quantify this activity. The ability of the *Bacillus* isolates to fix nitrogen was tested by inoculating them into nitrogen-free Jensen’s medium and incubating at 28°C for 7 days. Bacterial growth in this medium served as an indicator of their nitrogen-fixing potential, which was evaluated qualitatively based on the presence of visible growth. For measuring the production of Indole-3-Acetic Acid (IAA), bacterial cultures were grown in LB broth containing 0.1 g/L L-tryptophan and incubated at 28°C for 48 hours. After incubation, the cultures were centrifuged to separate the supernatant, which was then combined with Salkowski’s reagent in a 1:1 ratio. The development of a pink hue indicated the production of IAA, with the absorbance being measured at 530 nm using a spectrophotometer. A standard curve was used to calculate the concentration of IAA based on the absorbance readings.

#### Bacterial Extract Preparation for GC–MS Analysis

The bacterial isolate was cultivated for 48 hours in a 500 mL narrow-mouth Erlenmeyer flask (Borosil 4,980,024) containing 300 mL of nutrient broth (GM403 HiMedia). The incubation was maintained at 32 ± 2°C with continuous agitation to ensure proper aeration. After the incubation period, bacterial cells were separated by filtration using Whatman No.1 filter paper. The resulting filtrate was subjected to sequential extraction with ethyl acetate using a glass separating funnel (Borosil 6,400). The process allowed for the isolation of chemical metabolites from the bacterial cells. The collected extracts were then concentrated using a rotary evaporator (Model: RE100-Pro) under reduced pressure, maintaining a temperature range of 40-45°C. The concentrated extracts were stored at −20°C for preservation, following the method outlined by (Palanichamy et al. 2018). The ethyl acetate extract containing active components was dissolved back into ethyl acetate to achieve a final concentration of 1 mg/mL. This prepared solution was used for subsequent GC-MS analysis.

#### Gas Chromatography-Mass Spectrometry (GC-MS) Profiling of Bacterial Extract

The gas chromatography-mass spectrometry (GC-MS) analysis of the bacterial extract was conducted using a Perkin Elmer Clarus-680 gas chromatograph, equipped with a Clarus 600 mass spectrometer and a capillary column (30 m length, 0.25 mm internal diameter, 250 μm film thickness). The initial oven temperature was programmed at 60°C for 2 minutes, followed by a gradual ramp-up to 300°C at a rate of 10°C per minute, sustaining this final temperature for an additional 6 minutes. Helium served as the carrier gas, flowing at a constant rate of 1 mL/min. The transfer line and ion source temperatures were set at 240°C throughout the analysis. The entire analysis process took approximately 25 minutes. The spectral data were analyzed using TurboMass software (version 5.4.2), and the identification of chemical compounds was performed by comparing the acquired mass spectra with reference data from the NIST database (2008), as adapted from the work of (Gautam et al. 2016).

### Biochemical Characterization of Fungal Pathogen and Host Response

#### Isolation and Culture of Fungal Pathogen

The early blight-causing fungus *Alternaria solani* was isolated from infected tomato leaves and maintained as a pure culture. The collection of diseased leaves was done using polythene bags. Small segments (approximately 3 × 3 mm) were cut from the infected regions and surface sterilized with a 30% commercial bleach solution for 15 minutes, followed by three washes with sterile distilled water. Sterilized lesion sections were placed onto Potato Dextrose Agar (PDA) slants containing 2% agar (HIMEDIA, India) and incubated at 28°C in complete darkness. Within three days, fine thread-like hyphae emerged from the lesions. Hyphal tips were transferred to new PDA slants and incubated under identical conditions for 14 days to stimulate sporulation. The spores and conidia were analyzed using a compound microscope (Leica), with pure cultures being achieved through repeated sub-culturing and stored at 4°C.

#### Inoculation Method for Tomato Plants

The tomato cultivar PKM1 was selected for this investigation. The plants were divided into two groups: Co (control plants without microbial consortium application) and Bo (plants treated with a microbial consortium). 12-day-old *A. solani* culture served as the source of inoculum. Spore clusters were gently scraped from the culture plates and suspended in 20 mL of sterile distilled water. The spore suspension was then filtered through triple-layered cheesecloth. The concentration of spores was adjusted to 2 × 10⁵ spores/mL using a haemocytometer under a compound microscope. To promote uniform distribution over the leaf surfaces, 0.01% Tween 20 was added to the suspension as a surfactant. The inoculum was applied onto the leaves of three-week-old plants using an atomizer. The plants were kept under a light/dark cycle of 16 hours of light and 8 hours of darkness at 28°C for various periods (24, 48, and 72 hours).

#### Evaluation of Disease Progression

Disease development in the PKM1 tomato variety was monitored by measuring the necrotic area percentage on the leaves that had been inoculated. Disease severity levels were classified as follows: 0 = No lesion, 1 = Early signs of necrotic spots, 2 = Lesions covering under 50% of the leaf area, and 3 = Lesions covering over 50% of the leaf area.

#### Detached Leaf Infection Assay

Detached leaves measuring 2.5–3.5 cm from three-week-old PKM1 tomato plants were placed in Petri dishes lined with moistened filter paper. Fungal plugs, 5 mm in diameter, were taken from 10-day-old *A. solani* cultures using a cork borer. Each leaf was inoculated by positioning one fungal plug at its center. Petri dishes were incubated at 28°C under a 16-hour light/8-hour dark cycle. Disease progression was assessed based on lesion development and tissue breakdown caused by the pathogen, following methods adapted from (G. Liu et al. 2007). Experiments were performed in triplicate and repeated twice for consistency.

#### Visualization of Hydrogen Peroxide (H₂O₂) Localization Using DAB Staining

A modified 3,3ʹ-Diaminobenzidine (DAB) staining method, as described by (Thordal-Christensen et al. 1997), was employed to detect H₂O₂ buildup in the leaves. The primary leaves, following inoculation, were decolorized using an acetic acid-ethanol mixture (1:3) at room temperature for 24 hours. They were then treated with a 1 mg/mL DAB solution (Sigma Aldrich) at pH 3.8 and subjected to vacuum infiltration for 8 hours. Excess stain was removed using filter paper, and the leaves were mounted in 50% glycerol for observation under an Olympus BX-51 microscope.

#### Visualization of Phenolic Compounds Using Toluidine Blue

To detect phenolic compounds, toluidine blue staining was used. A 0.05% solution of toluidine blue was prepared in 0.1 M potassium phosphate buffer at pH 5.5. Leaves were decolorized with an acetic acid-ethanol solution (1:3) before being stained in toluidine blue for 6 hours. Stained samples were then mounted in 50% glycerol and inspected under a microscope, with phenolic presence indicated by a greenish-blue color, as outlined by (Borden and Higgins 2002).

#### Detection of Superoxide Dismutase (SOD) Activity Using NBT Staining

Superoxide dismutase (SOD) accumulation in the leaves was analyzed 24, 48, and 72 hours after inoculation. Leaves detached were decolorized in a solution of acetic acid and ethanol (1:3 by volume) and treated with a 0.5 mM nitro blue tetrazolium (NBT) solution at pH 7.4 for 12 hours. The leaves were then rinsed with deionized water to remove excess stain and mounted on slides with 50% glycerol for detailed observation under a microscope, following the method by Romero et al. (2008).

#### Coomassie Blue Staining for Cross-Linked Proteins

Cross-linked protein analysis was conducted on detached tomato leaves infected with *A. solani*, incubated for 24, 48, and 72 hours as previously described. After decolorizing with an acetic acid-ethanol mixture (1:3), the leaves were treated for 15 minutes with 1% SDS at 80°C. They were then stained using 0.1% Coomassie Blue for 10 to 30 minutes. Post-staining, the leaves were fixed with 2% glutaraldehyde and observed microscopically, according to methods described by Mellersh et al. (2002).

#### RNA Extraction and Semi-Quantitative RT-PCR Analysis

The leaves were collected from the middle third of the canopy from both microbial consortium-treated plants (BO) and control plants (CO). Six replicate plants were used for each group, and the collected leaves were pooled together. The pooled leaves were immediately flash-frozen in liquid nitrogen and stored at -80°C until further analysis. Total RNA was isolated from these frozen samples using the RNeasy Plant Mini Kit (Qiagen, Valencia, CA, USA), following the instructions provided by the manufacturer. RNA concentrations were determined by measuring absorbance at 260 nm using a spectrophotometer.

To eliminate any residual DNA contamination, the extracted RNA was treated with RNase-free DNase (MBI Fermentas, Germany). Synthesis of the first-strand cDNA was conducted using the RevertAid™ H Minus First Strand cDNA Synthesis Kit (MBI Fermentas, Germany) and an oligo dT primer, strictly adhering to the manufacturer’s protocol. The synthesized cDNA served as the template for subsequent semi-quantitative RT-PCR analysis, which was performed on a Thermo cycler (Flexi Gene-Techne, Cambridge, UK).

The PCR was initiated with an initial denaturation step at 94°C for 3 minutes. This was followed by 30 cycles, each consisting of denaturation at 94°C for 45 seconds, annealing at 64°C for 1 minute, and extension at 72°C for 1 minute. A final extension was carried out at 72°C for 3 minutes. To verify the absence of genomic DNA contamination, 200 ng of the DNase-treated RNA was used as a negative control in the PCR reactions.

The study targeted several defense-related genes, including Pto Kinase, PR2b, and Chi3, using the ubiquitin gene (Ubi3) as an internal control. The gene-specific primers were designed based on the work of (Khan, Mishra, and Nautiyal 2012) and synthesized by Eurofins Genomics (Bangalore, India). The sequences of the primers used for these genes are presented in Table 1.

The band intensities of the PCR products were measured using Scion Image Software (National Institute of Health, Maryland, USA). The values were normalized to the expression levels of the internal control gene (Ubi3) and expressed as relative percentages to provide a comparative analysis of gene expression levels.

### Metabolomic Analysis of Tomato Plants

#### Cultivation of Plant Material and Experimental Conditions

Tomato seeds of the PKM-1 variety were sterilized by immersing them in a 2% sodium hypochlorite solution for 5 minutes, followed by extensive rinsing with sterile distilled water. The cleaned seeds were dried on sterile filter paper and sown into earthen pots (dimensions 20 × 20 × 14 cm) filled with a sterile growth medium composed of red soil, cocopeat, perlite, and vermiculite, mixed in a 2:1:1:1 ratio by weight. Seedlings were cultivated in a controlled glasshouse environment, maintaining temperatures between 24°C and 28°C with relative humidity levels of 70-75%. At the early flowering stage, 60 days after germination, the treatment group (BO) received foliar applications of a microbial consortium, while the control group (CO) was sprayed with an equal volume of sterile water. Each treatment was replicated three times, with three biological replicates per group, labeled as CO1, CO2, CO3 for controls and BO1, BO2, BO3 for treated plants.

#### Preparation of Samples and Extraction of Metabolites

For metabolite analysis, upper leaves were harvested from both the treated and control plants seven days after the treatment application. The collected leaf samples were immediately frozen in liquid nitrogen and finely ground using a pre-chilled mortar and pestle. Approximately 2 grams of the powdered leaf tissue per sample was extracted using 5 mL of HPLC-grade methanol with sonication for 15 minutes. The extract was then centrifuged at 6000 × g for 15 minutes at 4°C to separate the supernatant. The obtained supernatants were dried using a rotary evaporator set at 50°C. The resulting dried extracts were reconstituted in 1 mL of HPLC-grade methanol, filtered through 0.22 µm syringe filters, and stored in dark glass vials at 4°C until further analysis.

#### LC-MS/MS-Based Metabolite Profiling

The metabolomic profiling was conducted using a Dionex Ultimate 3000 HPLC system integrated with a Q Exactive Plus Orbitrap mass spectrometer (Thermo Fisher Scientific, USA). Chromatographic separation utilized a Hypersil Gold C18 column (2.1 mm × 100 mm, 1.9 µm) maintained at 35°C. The mobile phase consisted of 0.05% formic acid in water (A) and 0.05% formic acid in acetonitrile (B), with a flow rate of 350 µL/min. The gradient elution began with 5% B, linearly ramped up to 95% B over 22 minutes, held at 95% B for 5 minutes, and then returned to 5% B over the final 4 minutes.

The mass spectrometer operated in both positive and negative ionization modes, with a scan range of 100 to 1500 m/z. Data-dependent MS/MS analysis was performed with a resolution of 35,000, using an AGC target of 1e6, a maximum injection time of 100 ms, and collision energies varying between 25-45 eV. Quality control samples were analyzed intermittently to ensure system stability and performance.

#### Data Analysis and Statistical Interpretation

Raw data from the mass spectrometry analysis were processed using Compound Discoverer 3.3 (Thermo Fisher Scientific, USA) for peak identification, alignment, and normalization. Metabolites were annotated using databases like mzCloud and ChemSpider, with further validation through KEGG pathway analysis. MetaboAnalyst 6.0 was employed for conducting multivariate statistical analyses, including Principal Component Analysis (PCA) and Orthogonal Partial Least Squares Discriminant Analysis (OPLS-DA). These analyses were used to distinguish between control and treated groups, identify potential biomarkers, and assess the treatment’s impact on metabolic pathways. The tools within MetaboAnalyst facilitated feature selection, pathway enrichment, and the creation of correlation networks, offering insights into the underlying metabolic responses.

## Acknowledgement

The authors would like to express their sincere gratitude to VIT University, Vellore, for providing the facilities and resources necessary to conduct this study.

## References

Agrios, George. 2004. “Plant Pathology: Fifth Edition.” Plant Pathology: Fifth Edition 9780080473789 (December). Academic Press: 1–922. doi:10.1016/C2009-0-02037-6.

Anzano, Attilio, Giuliano Bonanomi, Stefano Mazzoleni, and Virginia Lanzotti. 2021. “Plant Metabolomics in Biotic and Abiotic Stress: A Critical Overview.” Phytochemistry Reviews 2021 21:2 21 (2). Springer: 503–24. doi:10.1007/S11101-021-09786-W.

Bassem Yassin, Fatima, Mohammad Salal Aliwi, Saad Shakir Mahmood -, A Al-Maamori, Ahmad D Salman, Muntadher Al-Budeiri, J S Alkobaisy, E T Abdel Ghani, N A Mutlag, and A A Sh Lafi. 2021. “Effect of Vermicompost and Vermicompost Tea on the Growth and Yield of Broccoli and Some Soil Properties.” IOP Conference Series: Earth and Environmental Science 761 (1). IOP Publishing: 012008. doi:10.1088/1755-1315/761/1/012008.

Bonaterra, Anna, Esther Badosa, Núria Daranas, Jesús Francés, Gemma Roselló, and Emilio Montesinos. 2022. “Bacteria as Biological Control Agents of Plant Diseases.” Microorganisms 2022, Vol. 10, Page 1759 10 (9). Multidisciplinary Digital Publishing Institute: 1759. doi:10.3390/MICROORGANISMS10091759.

Borden, Stephanie, and Verna J. Higgins. 2002. “Hydrogen Peroxide Plays a Critical Role in the Defence Response of Tomato to Cladosporium Fulvum.” Physiological and Molecular Plant Pathology 61 (4). Academic Press: 227–36. doi:10.1006/PMPP.2002.0435.

Bozza, Desiree, Davide Barboni, Natasha Damiana Spadafora, Simona Felletti, Chiara De Luca, Chiara Nosengo, Greta Compagnin, Alberto Cavazzini, and Martina Catani. 2024. “Untargeted Metabolomics Approaches for the Characterization of Cereals and Their Derived Products by Means of Liquid Chromatography Coupled to High Resolution Mass Spectrometry.” Journal of Chromatography Open 6 (November). Elsevier: 100168. doi:10.1016/J.JCOA.2024.100168.

Brent, Keith J, and Derek W Hollomon. 2007. Fungicide Resistance in Plant Management: How Can It Be Managed? Fungicide Resistance Action Committee. www.frac.info.

Carvalhais, Lilia C., Paul G. Dennis, Dayakar V. Badri, Gene W. Tyson, Jorge M. Vivanco, and Peer M. Schenk. 2013. “Activation of the Jasmonic Acid Plant Defence Pathway Alters the Composition of Rhizosphere Bacterial Communities.” PLOS ONE 8 (2). Public Library of Science: e56457. doi:10.1371/JOURNAL.PONE.0056457.

Caulier, Simon, Annika Gillis, Gil Colau, Florent Licciardi, Maxime Liépin, Nicolas Desoignies, Pauline Modrie, Anne Legrève, Jacques Mahillon, and Claude Bragard. 2018. “Versatile Antagonistic Activities of Soil-Borne Bacillus Spp. and Pseudomonas Spp. against Phytophthora Infestans and Other Potato Pathogens.” Frontiers in Microbiology 9 (FEB). Frontiers Media S.A.: 304515. doi:10.3389/FMICB.2018.00143/BIBTEX.

Chaerani, Reni, and Roeland E. Voorrips. 2006. “Tomato Early Blight (Alternaria Solani): The Pathogen, Genetics, and Breeding for Resistance.” Journal of General Plant Pathology 72 (6). Springer: 335–47. doi:10.1007/S10327-006-0299-3/METRICS.

Chen, Shen, Xiaojing Wang, Yu Cheng, Hongsheng Gao, and Xuehao Chen. 2023. “A Review of Classification, Biosynthesis, Biological Activities and Potential Applications of Flavonoids.” Molecules 2023, Vol. 28, Page 4982 28 (13). Multidisciplinary Digital Publishing Institute: 4982. doi:10.3390/MOLECULES28134982.

Chen, Wumei, Zhansheng Wu, Changhao Liu, Ziyan Zhang, and Xiaochen Liu. 2023. “Biochar Combined with Bacillus Subtilis SL-44 as an Eco-Friendly Strategy to Improve Soil Fertility, Reduce Fusarium Wilt, and Promote Radish Growth.” Ecotoxicology and Environmental Safety 251 (February). Academic Press: 114509. doi:10.1016/J.ECOENV.2023.114509.

Choudhary, Devendra K., and Bhavdish N. Johri. 2009. “Interactions of Bacillus Spp. and Plants – With Special Reference to Induced Systemic Resistance (ISR).” Microbiological Research 164 (5). Urban & Fischer: 493–513. doi:10.1016/J.MICRES.2008.08.007.

Conrath, Uwe. 2006. “Systemic Acquired Resistance.” Plant Signaling & Behavior 1 (4). Taylor & Francis: 179–84. doi:10.4161/PSB.1.4.3221.

Dark, Adeeba, Vadim Demidchik, Siân L. Richards, Sergey Shabala, and Julia M. Davies. 2011. “Release of Extracellular Purines from Plant Roots and Effect on Ion Fluxes.” Plant Signaling & Behavior 6 (11). Taylor & Francis: 1855–57. doi:10.4161/PSB.6.11.17014.

Erb, Matthias, and Daniel J. Kliebenstein. 2020. “Plant Secondary Metabolites as Defenses, Regulators, and Primary Metabolites: The Blurred Functional Trichotomy.” Plant Physiology 184 (1). Oxford Academic: 39–52. doi:10.1104/PP.20.00433.

Fonseca, Sandra, Andrea Chini, Mats Hamberg, Bruce Adie, Andrea Porzel, Robert Kramell, Otto Miersch, Claus Wasternack, and Roberto Solano. 2009. “(+)-7-Iso-Jasmonoyl-L-Isoleucine Is the Endogenous Bioactive Jasmonate.” Nature Chemical Biology 5 (5): 344–50. doi:10.1038/nchembio.161.

Gao, Lei, Jinbiao Ma, Yonghong Liu, Yin Huang, Osama Abdalla Abdelshafy Mohamad, Hongchen Jiang, Dilfuza Egamberdieva, Wenjun Li, and Li Li. 2021. “Diversity and Biocontrol Potential of Cultivable Endophytic Bacteria Associated with Halophytes from the West Aral Sea Basin.” Microorganisms 9 (7). MDPI AG: 1448. doi:10.3390/MICROORGANISMS9071448/S1.

Gautam, Chandan Kumar, Mukund Madhav, Astha Sinha, and William Jabez Osborne. 2016. “VIT-CMJ2: Endophyte of Agaricus Bisporus in Production of Bioactive Compounds.” Iranian Journal of Biotechnology 14 (2). National Institute of Genetic Engineering and Biotechnology of Iran: 19–24. doi:10.15171/IJB.1287.

Gisi, Ulrich, and Helge Sierotzki. 2008. “Fungicide Modes of Action and Resistance in Downy Mildews.” European Journal of Plant Pathology 122 (1): 157–67. doi:10.1007/s10658-008-9290-5.

Gupta, Kartikey Kumar, Kamal Rai Aneja, and Deepanshu Rana. 2016. “Current Status of Cow Dung as a Bioresource for Sustainable Development.” Bioresources and Bioprocessing 3 (1). Springer Science and Business Media Deutschland GmbH: 1–11. doi:10.1186/S40643-016-0105-9/METRICS.

Harman, Gary E., Charles R. Howell, Ada Viterbo, Ilan Chet, and Matteo Lorito. 2004. “Trichoderma Species — Opportunistic, Avirulent Plant Symbionts.” Nature Reviews Microbiology 2004 2:1 2 (1). Nature Publishing Group: 43–56. doi:10.1038/nrmicro797.

Ishihara, Atsushi, Takahito Nakao, Yuko Mashimo, Masatoshi Murai, Naoya Ichimaru, Chihiro Tanaka, Hiromitsu Nakajima, Kyo Wakasa, and Hisashi Miyagawa. 2011. “Probing the Role of Tryptophan-Derived Secondary Metabolism in Defense Responses against Bipolaris Oryzae Infection in Rice Leaves by a Suicide Substrate of Tryptophan Decarboxylase.” Phytochemistry 72 (1). Pergamon: 7–13. doi:10.1016/J.PHYTOCHEM.2010.11.001.

Jaffar, Nur Sulastri, Roslina Jawan, and Khim Phin Chong. 2023. “The Potential of Lactic Acid Bacteria in Mediating the Control of Plant Diseases and Plant Growth Stimulation in Crop Production -A Mini Review.” Frontiers in Plant Science 13 (January). Frontiers Media S.A.: 1047945. doi:10.3389/FPLS.2022.1047945/BIBTEX.

Ji, Chao, Zhizhang Chen, Xuehua Kong, Zhiwen Xin, Fujin Sun, Jiahao Xing, Chunyu Li, Kun Li, Zengwen Liang, and Hui Cao. 2022. “Biocontrol and Plant Growth Promotion by Combined Bacillus Spp. Inoculation Affecting Pathogen and AMF Communities in the Wheat Rhizosphere at Low Salt Stress Conditions.” Frontiers in Plant Science 13 (December). Frontiers Media S.A.: 1043171. doi:10.3389/FPLS.2022.1043171/BIBTEX.

Jordan, Sean F., Eloise Nee, and Nick Lane. 2019. “Isoprenoids Enhance the Stability of Fatty Acid Membranes at the Emergence of Life Potentially Leading to an Early Lipid Divide.” Interface Focus 9 (6). Royal Society Publishing. doi:10.1098/RSFS.2019.0067.

Kadam, Rahul, Sangyeol Jo, Jonghwa Lee, Kamonwan Khanthong, Heewon Jang, and Jungyu Park. 2024. “A Review on the Anaerobic Co-Digestion of Livestock Manures in the Context of Sustainable Waste Management.” Energies 2024, Vol. 17, Page 546 17 (3). Multidisciplinary Digital Publishing Institute: 546. doi:10.3390/EN17030546.

Karun, Wilson, and Sathiavelu Arunachalam. 2023. “Phylogenomics Based Identification of Microbial Biocontrol Agent against Alternaria Solani.” Research Journal of Biotechnology 18 (9). World Research Association: 44–57. doi:10.25303/1809RJBT044057.

Khan, Noor, Aradhana Mishra, and Chandra Shekhar Nautiyal. 2012. “Paenibacillus Lentimorbus B-30488r Controls Early Blight Disease in Tomato by Inducing Host Resistance Associated Gene Expression and Inhibiting Alternaria Solani.” Biological Control 62 (2). Academic Press: 65–74. doi:10.1016/J.BIOCONTROL.2012.03.010.

Kim, Hyun Hwi, In Hye Jeong, Ja Shil Hyun, Byung Soo Kong, Ho Jin Kim, and Sung Jean Park. 2017. “Metabolomic Profiling of CSF in Multiple Sclerosis and Neuromyelitis Optica Spectrum Disorder by Nuclear Magnetic Resonance.” PLOS ONE 12 (7). Public Library of Science: e0181758. doi:10.1371/JOURNAL.PONE.0181758.

Kloepper, Joseph W., Choong Min Ryu, and Shouan Zhang. 2007. “Induced Systemic Resistance and Promotion of Plant Growth by Bacillus Spp.” 94 (11). The American Phytopathological Society : 1259–66. doi:10.1094/PHYTO.2004.94.11.1259.

Köhl, Jürgen, Rogier Kolnaar, and Willem J. Ravensberg. 2019. “Mode of Action of Microbial Biological Control Agents against Plant Diseases: Relevance beyond Efficacy.” Frontiers in Plant Science 10 (July). Frontiers Media S.A.: 454982. doi:10.3389/FPLS.2019.00845/BIBTEX.

Lamont, John R., Olivia Wilkins, Margaret Bywater-Ekegärd, and Donald L. Smith. 2017. “From Yogurt to Yield: Potential Applications of Lactic Acid Bacteria in Plant Production.” Soil Biology and Biochemistry 111 (August). Pergamon: 1–9. doi:10.1016/J.SOILBIO.2017.03.015.

Li, Fang, Liuyiqi Jiang, Shuming Pan, Shaowei Jiang, Yiwen Fan, Chao Jiang, Chengjin Gao, and Yuxin Leng. 2022. “Multi-Omic Profiling Reveals That Intra-Abdominal-Hypertension-Induced Intestinal Damage Can Be Prevented by Microbiome and Metabolic Modulations with 5-Hydroxyindoleacetic Acid as a Diagnostic Marker.” MSystems 7 (3). American Society for Microbiology. doi:10.1128/MSYSTEMS.01204-21/SUPPL_FILE/MSYSTEMS.01204-21-S0010.PDF.

Liu, Dongli, Rui Yan, Yansong Fu, Xiangjing Wang, Ji Zhang, and Wensheng Xiang. 2019. “Antifungal, Plant Growth-Promoting, and Genomic Properties of an Endophytic Actinobacterium Streptomyces Sp. NEAU-S7GS2.” Frontiers in Microbiology 10 (September). Frontiers Media S.A.: 471636. doi:10.3389/FMICB.2019.02077/BIBTEX.

Liu, Fei, Fangfang Wei, Lei Wang, Hui Liu, Xiaoping Zhu, and Yuancun Liang. 2010. “Riboflavin Activates Defense Responses in Tobacco and Induces Resistance against Phytophthora Parasitica and Ralstonia Solanacearum.” Physiological and Molecular Plant Pathology 74 (5–6). Academic Press: 330–36. doi:10.1016/J.PMPP.2010.05.002.

Liu, Guosheng, Regan Kennedy, David L. Greenshields, Gary Peng, Lily Forseille, Gopalan Selvaraj, and Yangdou Wei. 2007. “Detached and Attached Arabidopsis Leaf Assays Reveal Distinctive Defense Responses Against Hemibiotrophic Colletotrichum Spp.” 20 (10). The American Phytopathological Society : 1308–19. doi:10.1094/MPMI-20-10-1308.

Loon, L.C. van. 2000. “Induced System Resitance as a Mechanism Disease Suppression by Rhizobacteria.” *In* Mechanisms of Resistance to Plant Diseases. Dordrecht: Springer Netherlands, 521–74.

Monteiro Espíndola, Kaio Murilo, Roseane Guimarães Ferreira, Luis Eduardo Mosquera Narvaez, Amanda Caroline Rocha Silva Rosario, Agnes Hanna Machado Da Silva, Ana Gabrielle Bispo Silva, Ana Paula Oliveira Vieira, and Marta Chagas Monteiro. 2019. “Chemical and Pharmacological Aspects of Caffeic Acid and Its Activity in Hepatocarcinoma.” Frontiers in Oncology 9 (JUN). Frontiers Media S.A.: 467241. doi:10.3389/FONC.2019.00541/BIBTEX.

Munir, Nigarish, Chunzhen Cheng, Chaoshui Xia, Xuming Xu, Muhammad Azher Nawaz, Junaid Iftikhar, Yukun Chen, Yuling Lin, and Zhongxiong Lai. 2019. “RNA-Seq Analysis Reveals an Essential Role of Tyrosine Metabolism Pathway in Response to Root-Rot Infection in Gerbera Hybrida.” PLOS ONE 14 (10). Public Library of Science: e0223519. doi:10.1371/JOURNAL.PONE.0223519.

Nam, Nguyen Nhat, Hoang Dang Khoa Do, Kieu The Loan Trinh, and Nae Yoon Lee. 2023. “Metagenomics: An Effective Approach for Exploring Microbial Diversity and Functions.” Foods 12 (11). Multidisciplinary Digital Publishing Institute (MDPI). doi:10.3390/FOODS12112140.

Ongena, Marc, and Philippe Jacques. 2008. “Bacillus Lipopeptides: Versatile Weapons for Plant Disease Biocontrol.” Trends in Microbiology 16 (3). Elsevier Ltd: 115–25. doi:10.1016/J.TIM.2007.12.009/ASSET/C963C1FA-F689-453F-862D-4C9C5C5465AD/MAIN.ASSETS/GR1B3.SML.

Palanichamy, Prabha, Govindan Krishnamoorthy, Suganya Kannan, and Murugan Marudhamuthu. 2018. “Bioactive Potential of Secondary Metabolites Derived from Medicinal Plant Endophytes.” Egyptian Journal of Basic and Applied Sciences 5 (4). Taylor & Francis: 303–12. doi:10.1016/J.EJBAS.2018.07.002.

Pellegrini, Marika, Beatriz E Guerra-Sierra, Debasis Mitra, Asma Hasan, Baby Tabassum, Mohammad Hashim, and Nagma Khan. 2024. “Role of Plant Growth Promoting Rhizobacteria (PGPR) as a Plant Growth Enhancer for Sustainable Agriculture: A Review.” Bacteria 2024, Vol. 3, Pages 59-75 3 (2). Multidisciplinary Digital Publishing Institute: 59–75. doi:10.3390/BACTERIA3020005.

Pieterse, Corné M.J., Christos Zamioudis, Roeland L. Berendsen, David M. Weller, Saskia C.M. Van Wees, and Peter A.H.M. Bakker. 2014. “Induced Systemic Resistance by Beneficial Microbes.” Annual Review of Phytopathology 52 (Volume 52, 2014). Annual Reviews Inc.: 347–75. doi:10.1146/ANNUREV-PHYTO-082712-102340/CITE/REFWORKS.

Rahim, Md Abdur, Xuebin Zhang, and Nicola Busatto. 2023. “Editorial: Phenylpropanoid Biosynthesis in Plants.” Frontiers in Plant Science 14 (June). Frontiers Media SA: 1230664. doi:10.3389/FPLS.2023.1230664/BIBTEX.

Shi, Huimin, Xiaoxia Zhu, Lanxiang Lu, and Jianren Ye. 2024. “Effect of Microbial Inoculants Endowed with Multifarious Plant Growth-Promoting Traits on Grape Growth and Fruit Quality under Organic Fertilization Scenarios.” Agronomy 14 (3). Multidisciplinary Digital Publishing Institute (MDPI): 491. doi:10.3390/AGRONOMY14030491/S1.

Shleeva, Margarita O., Daria A. Kondratieva, and Arseny S. Kaprelyants. 2023. “Bacillus Licheniformis: A Producer of Antimicrobial Substances, Including Antimycobacterials, Which Are Feasible for Medical Applications.” Pharmaceutics 2023, Vol. 15, Page 1893 15 (7). Multidisciplinary Digital Publishing Institute: 1893. doi:10.3390/PHARMACEUTICS15071893.

Stein, Torsten. 2005. “Bacillus Subtilis Antibiotics: Structures, Syntheses and Specific Functions.” Molecular Microbiology 56 (4). John Wiley & Sons, Ltd: 845–57. doi:10.1111/J.1365-2958.2005.04587.X.

Thordal-Christensen, Hans, Ziguo Zhang, Yangdou Wei, and David B. Collinge. 1997. “Subcellular Localization of H2O2 in Plants. H2O2 Accumulation in Papillae and Hypersensitive Response during the Barley-Powdery Mildew Interaction.” The Plant Journal 11 (6). Wiley: 1187–94. doi:10.1046/J.1365-313X.1997.11061187.X.

Vasques, Natalia Caetano, Marco Antonio Nogueira, and Mariangela Hungria. 2024. “Increasing Application of Multifunctional Bacillus for Biocontrol of Pests and Diseases and Plant Growth Promotion: Lessons from Brazil.” Agronomy 2024, Vol. 14, Page 1654 14 (8). Multidisciplinary Digital Publishing Institute: 1654. doi:10.3390/AGRONOMY14081654.

Vishwakarma, Kanchan, Nitin Kumar, Chitrakshi Shandilya, Swati Mohapatra, Sahil Bhayana, and Ajit Varma. 2020. “Revisiting Plant–Microbe Interactions and Microbial Consortia Application for Enhancing Sustainable Agriculture: A Review.” Frontiers in Microbiology 11 (December). Frontiers Media S.A.: 560406. doi:10.3389/FMICB.2020.560406/BIBTEX.

Vogt, Thomas. 2010. “Phenylpropanoid Biosynthesis.” Molecular Plant 3 (1). Elsevier: 2–20. doi:10.1093/MP/SSP106.

Wang, Xin, Tao Liu, Haixin Song, Shaoyang Cui, Gang Liu, Andrea Christoforou, Patrick Flaherty, Xun Luo, Lisa Wood, and Qing Mei Wang. 2020. “Targeted Metabolomic Profiling Reveals Association Between Altered Amino Acids and Poor Functional Recovery After Stroke.” Frontiers in Neurology 10 (January): 1–12. doi:10.3389/fneur.2019.01425.

Xie, Yanjie, Daokun Xu, Weiti Cui, and Wenbiao Shen. 2012. “Mutation of Arabidopsis HY1 Causes UV-C Hypersensitivity by Impairing Carotenoid and Flavonoid Biosynthesis and the down-Regulation of Antioxidant Defence.” Journal of Experimental Botany 63 (10). Oxford Academic: 3869–83. doi:10.1093/JXB/ERS078.

Yang, Jing, Guihua Duan, Chunqin Li, Lin Liu, Guangyu Han, Yaling Zhang, and Changmi Wang. 2019. “The Crosstalks Between Jasmonic Acid and Other Plant Hormone Signaling Highlight the Involvement of Jasmonic Acid as a Core Component in Plant Response to Biotic and Abiotic Stresses.” Frontiers in Plant Science 10 (October). Frontiers Media S.A.: 458580. doi:10.3389/FPLS.2019.01349/BIBTEX.

Yao, Tao, Kai Feng, Meng Xie, Jaime Barros, Timothy J. Tschaplinski, Gerald A. Tuskan, Wellington Muchero, and Jin Gui Chen. 2021. “Phylogenetic Occurrence of the Phenylpropanoid Pathway and Lignin Biosynthesis in Plants.” Frontiers in Plant Science 12 (August). Frontiers Media S.A.: 704697. doi:10.3389/FPLS.2021.704697/BIBTEX.

Zahoor, Sidra, Rabia Naz, Rumana Keyani, Thomas H. Roberts, Muhammad N. Hassan, Humaira Yasmin, Asia Nosheen, and Saira Farman. 2022. “Rhizosphere Bacteria Associated with Chenopodium Quinoa Promote Resistance to Alternaria Alternata in Tomato.” Scientific Reports 2022 12:1 12 (1). Nature Publishing Group: 1–16. doi:10.1038/s41598-022-21857-2.

Zhang, Dai, Shuiqing Yu, Yiqing Yang, Jinglin Zhang, Dongmei Zhao, Yang Pan, Shasha Fan, Zhihui Yang, and Jiehua Zhu. 2020. “Antifungal Effects of Volatiles Produced by Bacillus Subtilis Against Alternaria Solani in Potato.” Frontiers in Microbiology 11 (June). Frontiers Media S.A.: 545349. doi:10.3389/FMICB.2020.01196/BIBTEX.

Zhang, Shuning, Litao Sun, Yu Wang, Kai Fan, Qingshan Xu, Yusheng Li, Qingping Ma, Jiguo Wang, Wanming Ren, and Zhaotang Ding. 2020. “Cow Manure Application Effectively Regulates the Soil Bacterial Community in Tea Plantation.” BMC Microbiology 20 (1). BMC. doi:10.1186/S12866-020-01871-Y.

